# The APETALA2 transcription factor LsAP2 regulates seed shape in lettuce

**DOI:** 10.1101/2020.11.05.370684

**Authors:** Chen Luo, Shenglin Wang, Kang Ning, Zijing Chen, Yixin Wang, Jingjing Yang, Meixia Qi, Qian Wang

## Abstract

Seeds are major vehicles of propagation and dispersal in plants. A number of transcription factors, including APETALA2 (AP2), play crucial roles during the seed development process in various plant species. However, genes essential for seed development and the regulatory networks during seed development remain unclear in lettuce. Here, we identified a lettuce *AP2* (*LsAP2*) gene that was highly expressed at the early stages of seed development. *LsAP2* knockout plants obtained by the CRISPR/Cas9 system were used to explore the biological function of *LsAP2*. Compared with wild-type, the seeds of the *Lsap2* mutant plants had increased length and decreased width, and developed an extended tip at the seed top. After further investigating the seed structural characteristics of *Lsap2* mutant plants, we proposed a new function of *LsAP2* in seed dispersal. Moreover, we identified several interactors of LsAP2. Our results showed that LsAP2 directly interacted with the lettuce homolog of BREVIPEDICELLUS (LsBP) and promoted the expression of *LsBP*. Transcriptome analysis revealed that LsAP2 might also be involved in brassinosteroid biosynthesis and signaling pathways. Taken together, our data indicate that LsAP2 has a significant function in regulating seed shape in lettuce.

## Introduction

In angiosperms, a mature seed has three major components, embryo, endosperm, and seed coat. The seeds that store essential substances and contain genetic materials are the most important harvested organ in many plant species. Moreover, seed size and seed shape are crucial for plants to adapt during evolution, and seed size is also an important agronomic trait in crop domestication (Moles *et al.*, 2005).

Lettuce (*Lactuca sativa* L.) belongs to the Asteraceae family and is one of the most important leafy vegetables worldwide. The seed structure of lettuce is different from the model plant *Arabidopsis* as the lettuce seed is actually an achene fruit. Therefore, the achene development integrates the processes of fruit and seed development. Achene has four major components, pericarp, integuments, endosperm and embryo. The pericarp is derived from the ovary walls, and the pericarp and integuments are tightly integrated to constitute the seed coat (Steinbrecher and Leubner-Metzger, 2017). Lettuce seeds are the vehicles of propagation and dispersal, and many traits have changed dramatically during lettuce domestication, including seed size, seed shattering, and seed dispersal (Zhang *et al.*, 2017). A previous study showed that some lettuce MADS-box genes were highly expressed in seeds, and these genes may be involved in seed development (Ning *et al.*, 2019). However, few studies have performed the gene function analysis and explored the regulatory networks associated with seed development in lettuce.

Seed development is initiated by double fertilization (Jiang and Köhler, 2012). The zygote, the fertilized central cell and the maternal integuments develop together into a complete seed. Thus, the seed growth is coordinately controlled by the growth of maternal and zygotic tissues (Sreenivasulu and Wobus, 2013). Seed development is a complicated process modulated by complex networks. Recent studies have identified various pathways that control seed development through maternal tissues, including the ubiquitin-proteasome pathway, G-protein signaling pathway, mitogen-activated protein kinase signaling pathway, phytohormone signaling and homeostasis pathway, as well as various kinds of transcriptional regulatory factors (Li and Li, 2015; Li *et al.*, 2019). Many genes involved in the seed development process have been studied extensively, and many key regulators of seed size and seed shape have been identified (Sreenivasulu and Wobus, 2013; Li *et al.*, 2019).

*APETALA2* (*AP2*) gene belongs to the euAP2 lineage, a subgroup of the large AP2/ERF gene family. Genes in the euAP2 lineage encode two highly conserved AP2 domains and contain a microRNA172 (miR172) binding site (Kim *et al.*, 2006; Zumajo-Cardona and Pabon-Mora, 2016). *AP2* gene encodes a transcription factor, and its roles in specifying floral organ identity and regulating the expression of floral homeotic genes are well known (Kunst *et al.*, 1989; Bowman *et al.*, 1991; Coen and Meyerowitz, 1991; Drews *et al.*, 1991). Meanwhile, miR172 regulates the expression of *AP2* gene primarily through translational inhibition (Aukerman and Sakai, 2003; Chen, 2004). The *AP2* gene was first isolated in *Arabidopsis*, and *AP2* activity was shown to be required during seed development (Jofuku *et al.*, 1994). A mutation in the *AP2* gene (*ap2-1*) in *Arabidopsis* had a pleiotropic effect on seed shape, which changed from the normal oblong shape to a number of aberrant shapes (Leon-Kloosterziel *et al.*, 1994). AP2 was shown to act as an important negative regulator of seed development because *ap2* mutants produce large seeds. AP2 acts maternally to determine seed weight and seed yield by coordinating the growth of embryo, endosperm, and maternal integuments (Jofuku *et al.*, 2005; Ohto *et al.*, 2005; Ohto *et al.*, 2009). Petunia *AP2A*, an ortholog of *Arabidopsis AP2*, is also expressed in seeds, and it was found to restore the structural defect of seed epidermal cells in *Arabidopsis ap2-1* mutant (Maes *et al.*, 2001). Grain yield is an important agronomic trait in crop domestication. In rice, *SUPERNUMERARY BRACT* (*SNB*) encodes an AP2-like transcription factor (Lee *et al.*, 2007), and *SUPPRESSION OF SHATTERING1* (*SSH1*) was recently found to be an allele of *SNB*. The rice *ssh1* mutant shows reduced shattering and has larger seeds and higher grain weight (Jiang *et al.*, 2019). A genome-wide association study of grain length and grain width in rice also identified *SNB* as a novel gene that negatively regulates grain size. The *SNB* knockout plants show increased grain length, width, and weight (Ma *et al.*, 2019).

Nowadays, the CRISPR/Cas systems have received widespread attention because of its high specificity and efficiency in editing the genomes of humans, animals, and plants (Cong *et al.*, 2013; Shan *et al.*, 2013; Kumar and Jain, 2015), and scientists have applied the CRISPR/Cas9 system to precisely modify crop traits and accelerate the breeding process (Botella, 2019; Chen *et al.*, 2019). In this study, we identified a lettuce *AP2* (*LsAP2*) gene that was highly expressed at the early stages of seed development. Then, we used the CRISPR/Cas9 system to edit *LsAP2* and found that LsAP2 regulates lettuce seed shape. We also showed LsAP2-related regulatory networks during seed development in lettuce. Transcriptome analysis identified the differentially expressed genes (DEGs) between WT and *Lsap2* mutant plants. Moreover, LsAP2 might regulate seed shape through its effects on the brassinosteroid (BR) biosynthesis and signaling pathways.

## Materials and methods

### Plant materials and growth conditions

Lettuce cultivar S39 was used as wild-type (WT) in this study. The lettuce seeds were provided by Dr. Han from Beijing University of Agriculture (Han *et al.*, 2016). The lettuce plants were grown in growth chambers under a photoperiod of 16-h light (200 μmol m^−2^ s^−1^) at 25°C and 8-h dark at 18°C. When the fifth true leaf was fully expanded, the lettuce plants were transplanted to a greenhouse in the Science Park of China Agricultural University under standard greenhouse conditions.

### Phylogenetic analysis

Lettuce *AP2* genes were identified using BLASTp from the lettuce genome (Reyes-Chin-Wo *et al.*, 2017). Lettuce sequences were downloaded from the CoGe genome database (https://genomevolution.org/coge/). *Arabidopsis* sequences were downloaded from The *Arabidopsis* Information Resource (TAIR; https://www.arabidopsis.org/index.jsp). Petunia sequences were downloaded from the Sol Genomics Network (SGN; https://solgenomics.net/organism/Petunia_axillaris/genome). Sequences of tomato and rice were downloaded from the Phytozome genome database (https://phytozome.jgi.doe.gov/pz/portal.html). Sequences of maize and wheat were downloaded from the National Center for Biotechnology Information (NCBI; https://www.ncbi.nlm.nih.gov). The amino acid sequences were aligned using ClustalW (Larkin *et al.*, 2007). The phylogenetic tree was constructed in MEGA 5 program using the neighbor-joining method with 1000 bootstrap replicates (Tamura *et al.*, 2011). The accession numbers of the used sequences are listed in Supplementary Table S1.

### RNA extraction, cDNA synthesis, and qRT-PCR

Total RNA was extracted from different lettuce tissues using the Quick RNA isolation Kit (Huayueyang, China). The cDNAs were synthesized using the FastKing-RT SuperMix (Tiangen, China) according to the manufacturer’s protocol. Subsequently, quantitative real-time PCR (qRT-PCR) assays were performed in the QuantStudio 6 Flex Real-Time PCR System (Applied Biosystems, USA) using TB Green Premix Ex Taq II (Takara, Japan). The qRT-PCR assays were performed with technical triplicates of three biological replicates. The expression data were normalized using *LsPP2A-1* and *LsTIP41* as reference genes (Sgamma *et al.*, 2016). Relative expression levels were calculated using the 2^−ΔΔCT^ method (Livak and Schmittgen, 2001). Primers used for qRT-PCR assays are listed in Supplementary Table S2.

### Construction of the lettuce transformation vectors

To generate the *Lsap2* mutants, pKSE401-*LsAP2* was constructed using the previously described method (Xing *et al.*, 2014). The single-guide RNAs (sgRNAs) were designed using the Cas-Designer Tool (Park *et al.*, 2015; Liu *et al.*, 2017). Cas-OFFinder was used to predict the potential off-target sites of Cas9 RNA-guided endonucleases (Bae *et al.*, 2014; Liu *et al.*, 2017). The β-glucuronidase (*GUS*) reporter gene under the control of *LsAP2* promoter was used to generate the *pLsAP2:GUS* construct. Briefly, the 2402-bp genomic fragment upstream of the *LsAP2* start codon was amplified by PCR using genomic DNA as the template. Then, the fragment was inserted into the *Hind*III-*Bam*HI sites of the pBI121 vector to drive the *GUS* reporter gene. Primers used to generate the above constructs are listed in Supplementary Table S2.

### *Agrobacterium*-mediated transformation of lettuce

Lettuce transformation procedure was improved from the previously described method (Chen *et al.*, 2017). The detailed steps are as follows. Seeds of lettuce cultivar S39 were sterilized with 1.0% sodium hypochlorite for 10 min and rinsed with sterile water, then sown on MS medium (MS supplemented with 3% sucrose and 0.7% agar). The plates were incubated under a photoperiod of 16-h light (200 μmol m^−2^ s^−1^) at 25°C and 8-h dark at 18°C for about 5 days. When the cotyledons were expanded, they were cut and incubated with *Agrobacterium* GV3101 carrying the desired plasmid for 10 min in MS liquid medium (1/2 MS supplemented with 3% sucrose and 200 μmol/L acetosyringone). After incubation, the cotyledons were transferred to sterile filter papers to remove excess *Agrobacterium*. Then, the cotyledons were placed on MS co-cultivation medium (MS supplemented with 3% sucrose, 0.7% agar, 0.1 mg/L 6-benzylaminopurine, 0.05 mg/L α-naphthaleneacetic acid, and 200 μmol/L acetosyringone) and incubated at 25°C in the dark for 2 days. Subsequently, explants were transferred to MS selection medium (MS supplemented with 3% sucrose, 0.7% agar, 0.1 mg/L 6-benzylaminopurine, 0.05 mg/L α-naphthaleneacetic acid, 300 mg/L timentin, and 40 mg/L kanamycin monosulfate) and incubated under a cycle of 16-h light (200 μmol m^−2^ s^−1^) at 25°C and 8-h dark at 18°C. About two weeks later, regenerated shoots grew on the calli, and they were cut into MS shoot-inducing medium (MS supplemented with 3% sucrose, 0.7% agar, and 300 mg/L timentin). After two weeks, the shoots were subcultured on fresh MS shoot-inducing medium. When the shoots were 2-cm long, they were transferred to MS rooting medium (1/2 MS supplemented with 3% sucrose, 0.7% agar, 0.02 mg/L α-naphthaleneacetic acid, and 200 mg/L timentin). When the shoots and roots of the regenerated plants were well developed, they were transferred to the soil under standard growth conditions.

### GUS staining

Tissues from the *pLsAP2:GUS* plants were used for GUS staining assays. First, flowers and seeds at different developmental stages were collected and immersed in GUS staining buffer (containing 1 mM 5-bromo-4-chloro-3-indolyl-β-D-glucuronide). Next, the samples were vacuum infiltrated for 10 min, and then stained in a 37°C incubator overnight. After incubation, the staining buffer was removed, and samples were washed with 70% ethanol for several times to remove the pigments. Finally, the samples were immersed in fresh 70% ethanol before observation. The samples were observed using a Leica stereomicroscope (Leica S8 APO, Germany).

### Phenotypic analysis

Plant materials were photographed with a digital camera (Nikon D7000, Japan) and a stereomicroscope (Leica S8 APO, Germany). To evaluate seed size, the images were imported into ImageJ software to measure the seed length and seed width. The measured seed length and seed width were used to calculate the ratio of seed length to width. To obtain the scanning electron micrographs, the seeds were gold plated and observed using a scanning electron microscope (Hitachi S-3400N, Japan). To evaluate cell size, the images were imported into ImageJ software to measure the cell length and cell width. Mature seeds were evaluated for calculating 1,000-grain weight using an electronic analytical balance (METTLER TOLEDO, USA). For seed germination tests, 20 seeds were tested for each line with three replicates, then the seeds were moistened with water and incubated at 25°C for two days to calculate the germination rates. To measure the seed beak diameter, beaks were photographed using a microscope (Olympus IX71, Japan), then the images were imported into ImageJ software to calculate the beak diameter. To evaluate seed dispersal, the breaking tensile strength (BTS) between beak and seed was measured with a digital force gauge (SHSIWI, China) at the harvest stage.

### Yeast two-hybrid assays

The full-length coding sequence (CDS) of *LsAP2* was cloned into pGADT7 vector. The full-length CDSs of *LsAS1*, *LsAS2*, *LsBP*, *LsKAN1*, and *LsKAN2* were amplified and cloned into pGBKT7 vector. Yeast two-hybrid assays were performed as described previously (Ding *et al.*, 2015). Briefly, bait plasmids and prey plasmids were co-transformed into yeast strain AH109 according to the manufacturer’s protocol of GAL4-based yeast two-hybrid system (Clontech, USA). Then, yeast transformants were grown on SD/-Leu-Trp plates for growth of the colonies and grown on SD/-Leu/-Trp/-His/-Ade plates for protein interaction selection. Primers used for yeast two-hybrid assays are listed in Supplementary Table S2.

### Firefly luciferase complementation imaging (LCI) assays

The full-length CDSs without the stop codon of *LsAS2*, *LsBP*, *LsKAN1*, and *LsKAN2* were cloned into the *35S:nLUC* vector. The full-length CDS of *LsAP2* was cloned into the *35S:cLUC* vector. Then, the above constructs were transformed into *Agrobacterium* strain GV3101. LCI assays were performed using the previously described method (Chen *et al.*, 2008). Briefly, different combinations were co-infiltrated into tobacco leaves using *Agrobacterium*-mediated transient transformation. After infiltration, the plants were grown under a 16-h light/8-h dark cycle for 2 days. Before imaging, the abaxial sides of the tobacco leaves were sprayed with luciferin, and the leaves were kept in the dark for 5 min. Then a cooled CCD camera (Roper 1300B, USA) was used to capture the fluorescence signal with 10 min exposures. Primers used for LCI assays are listed in Supplementary Table S2.

### Dual-luciferase reporter assays

The *LsBP* promoter (1595-bp genomic fragment upstream the start codon) was cloned and inserted into the pGreenII 0800-LUC vector as the reporter. The full-length CDSs of *LsAP2* and *LsBP* were inserted into the pGreenII 62-SK vector as effectors. The empty vector pGreenII 62-SK was used as the negative control. Dual-luciferase reporter assays were performed as described previously (Hellens *et al.*, 2005). Briefly, *Agrobacterium* strain GV3101 carrying the reporter and effector constructs were infiltrated into tobacco leaves. After infiltration, the plants were grown under a 16-h light/8-h dark cycle for 2 days. Luciferase (LUC) and renilla luciferase (REN) activities were measured by a multiscan spectrum (Molecular Devices SpectraMax® i3x, USA) using the Dual-Luciferase Reporter Assay Kit (Promega, USA) according to the manufacturer’s protocol. The ratio of LUC to REN was calculated as the final transcriptional activity. Primers used for dual-luciferase reporter assays are listed in Supplementary Table S2.

### Transcriptome analysis

Total RNA was extracted from the seeds of WT and *Lsap2* mutant plants on the day of pollination, with three biological replicates. Paired-end libraries were constructed and sequenced using an Illumina NovaSeq 6000 platform at Biomarker Technologies (BioMarker, China). Analyses of the transcriptome data were performed on the BMKCloud platform (www.biocloud.net). Clean reads were mapped to the lettuce reference genome (Reyes-Chin-Wo *et al.*, 2017) using the HISAT2 software (Kim *et al.*, 2015). Differential expression analysis between WT and *Lsap2* mutant plants (fold change ≥ 2, FDR < 0.01) was performed using the DESeq (Anders and Huber, 2010). Gene Ontology (GO) enrichment analysis and Kyoto Encyclopedia of Genes and Genomes (KEGG) analysis of the DEGs were performed using GOseq R packages and KOBAS software (Mao *et al.*, 2005; Young *et al.*, 2010).

### Measurement of endogenous hormones

Extraction and purification of the endogenous hormones were performed as described previously (Wang *et al.*, 2012). Briefly, about 0.2 g of the fresh seed samples from WT and *Lsap2* mutant plants were collected and homogenized in 4 mL of 80% methanol (containing antioxidant), with three biological replicates. After incubation at 4°C for 48 h, the extract was centrifuged at 5000 rpm for 10 min at 4°C. Then, the supernatant was passed through the extraction cartridge, and the phytohormone fraction was eluted with methanol and then ether. Next, the eluate was dried down and dissolved in phosphate-buffered saline (PBS) buffer. Quantification of auxin (IAA) and BR were measured by enzyme-linked immunosorbent assay (ELISA) following the described protocol (Zhao *et al.*, 2006) at the Center of Crop Chemical Control, China Agricultural University.

## Results

### Identification of *AP2* genes in lettuce

To understand the seed development process in lettuce and identify the key regulators during lettuce seed development, we continuously observed the developing seeds after pollination. We found that the seed morphology was fully developed within ten days after pollination (Fig. 1A). The seeds grew longer and wider in the first three days. Then, the seeds began to expand, and obvious longitudinal ribs were observed on the seed surface. One week later, the seed coat became darker, and the longitudinal ribs were more pronounced. Ultimately, the seeds were ready to ripen.

**Fig. 1.**
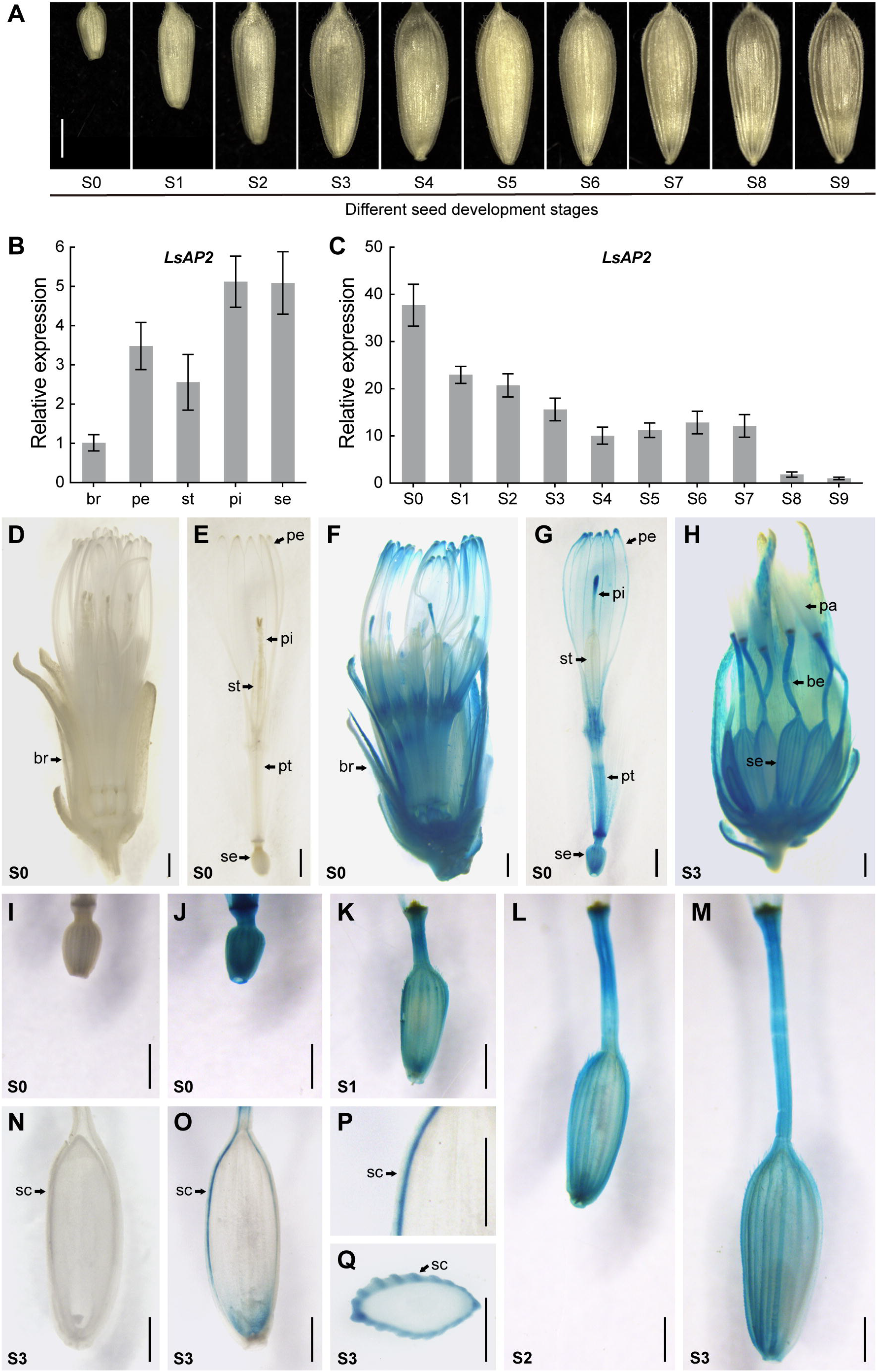
Expression pattern of *LsAP2*. (A) Different seed development stages. S0, the day of pollination; S1-S9, 1-9 days after pollination (DAP), respectively. (B) Expression of *LsAP2* in floral organs. br, bract; pe, petal; st, stamen; pi, pistil; se, seed. Values are means ± SD (*n* = 3). (C) Expression of *LsAP2* at different seed development stages. Values are means ± SD (*n* = 3). The expression data were normalized using *LsPP2A-1* and *LsTIP41* as reference genes. (D) The capitulum of WT at S0 stage. (E) A floret in D. pt, petal tube. (F) The capitulum of *pLsAP2:GUS* plant at S0 stage. (G) A floret in F. (H) The seed head of *pLsAP2:GUS* plant at S3 stage. pa, pappus; be, beak. (I) The seed of WT at S0 stage. (J-M) The seeds of *pLsAP2:GUS* plants at S0, S1, S2, and S3 stages, respectively. (N) Longitudinal section of the seed from WT. sc, seed coat (pericarp and integuments). (O) Longitudinal section of the seed from *pLsAP2:GUS* plant. (P) Magnified image of the longitudinal section in O. (Q) Transverse section of the seed from *pLsAP2:GUS* plant. Scale bars: 1 mm.

Previous studies have identified many transcription factors as important regulators of seed development (Li *et al.*, 2019), and *AP2* gene has been shown to play crucial roles in regulating seed development in many species. To investigate whether lettuce *AP2* gene plays a role in seed development, we performed BLAST searches at the lettuce genome database and identified three lettuce *AP2* genes as the candidate genes. The phylogenetic tree revealed that LsAP2 was more closely related to *Arabidopsis* AP2 than the other two lettuce AP2 homologs, namely LsAP2-like1 and LsAP2-like2 (Supplementary Fig. S1).

To functionally characterize the lettuce *AP2* genes during seed development, we examined their expression patterns by analyzing the FPKM values from a transcriptome dataset. Strikingly, *LsAP2* was the most highly expressed gene among the three lettuce *AP2* genes at different seed development stages, and the expression of *LsAP2* gradually decreased between successive stages (Supplementary Table S3). According to the homology and expression analysis, we speculated that of the three lettuce *AP2* genes, *LsAP2* might be the one that plays a major role during lettuce seed development. Therefore, we chose *LsAP2* for further characterization in this study.

### Expression pattern of *LsAP2*

To detect the expression pattern of *LsAP2* further, we analyzed its expression in different lettuce tissues by qRT-PCR assays. We found that *LsAP2* was expressed in all lettuce tissues, and the expression level of *LsAP2* in flower was many times higher than in other tissues (Supplementary Fig. S2). Further, we examined the expression of *LsAP2* in floral organs. *LsAP2* was highly expressed in petal, pistil, and seed, while *LsAP2* displayed lower expression level in bract and stamen (Fig. 1B). To explore when *LsAP2* was expressed during seed development, we then analyzed the expression of *LsAP2* in seeds at different developmental stages. Significantly, *LsAP2* transcripts were strongly accumulated at the early stages of seed development, and its expression decreased gradually as the seed developed (Fig. 1C). These results confirmed that *LsAP2* was expressed in seeds, especially at the early stages of seed development.

Further, we analyzed the spatial and temporal expression pattern of *LsAP2* in flower using the *GUS* reporter gene under the control of *LsAP2* promoter. The observed GUS activities were generally consistent with the endogenous *LsAP2* mRNA level determined by qRT-PCR. GUS staining showed that *LsAP2* was mainly expressed in petal, pistil, and seed, while weaker or even no GUS signal was observed in bract and stamen (Fig. 1F-H). Then, we analyzed the expression of *LsAP2* at different seed development stages. Strong GUS activities were also detected in seeds at the corresponding stages (Fig. 1J-M). Further, we observed different sections of the stained seeds, and only strong GUS signal was found in the seed coat (Fig. 1O-Q).

### CRISPR/Cas9-mediated editing of *LsAP2* alters seed shape

To investigate the biological function of *LsAP2* gene, we generated *LsAP2* knockout plants using the CRISPR/Cas9 system (Fig. 2A). Genotyping analysis of the ten T0 transgenic plants identified six plants with mutations in the target sites. There were significant differences in the editing efficiency of the two targets; Target 1 was mutated in six plants, whereas Target 2 was mutated in only one plant (Supplementary Figs. S3, S4). Therefore, it is crucial to assess the on-target efficiency when design sgRNAs. There were two types of mutations among the mutant plants; two plants were homozygous, and four plants were biallelic that had two mutant alleles (Supplementary Table S4). We also assessed the potential off-target sites using Cas-OFFinder, and no mutation was detected in the predicted off-target sites (Supplementary Table S5). Through the self-pollination of lettuce, homozygous and non-transgenic T1 generation plants were obtained and confirmed by PCR genotyping and sequencing. Three mutant lines were chosen for further exploration (Fig. 2B, C).

**Fig. 2.**
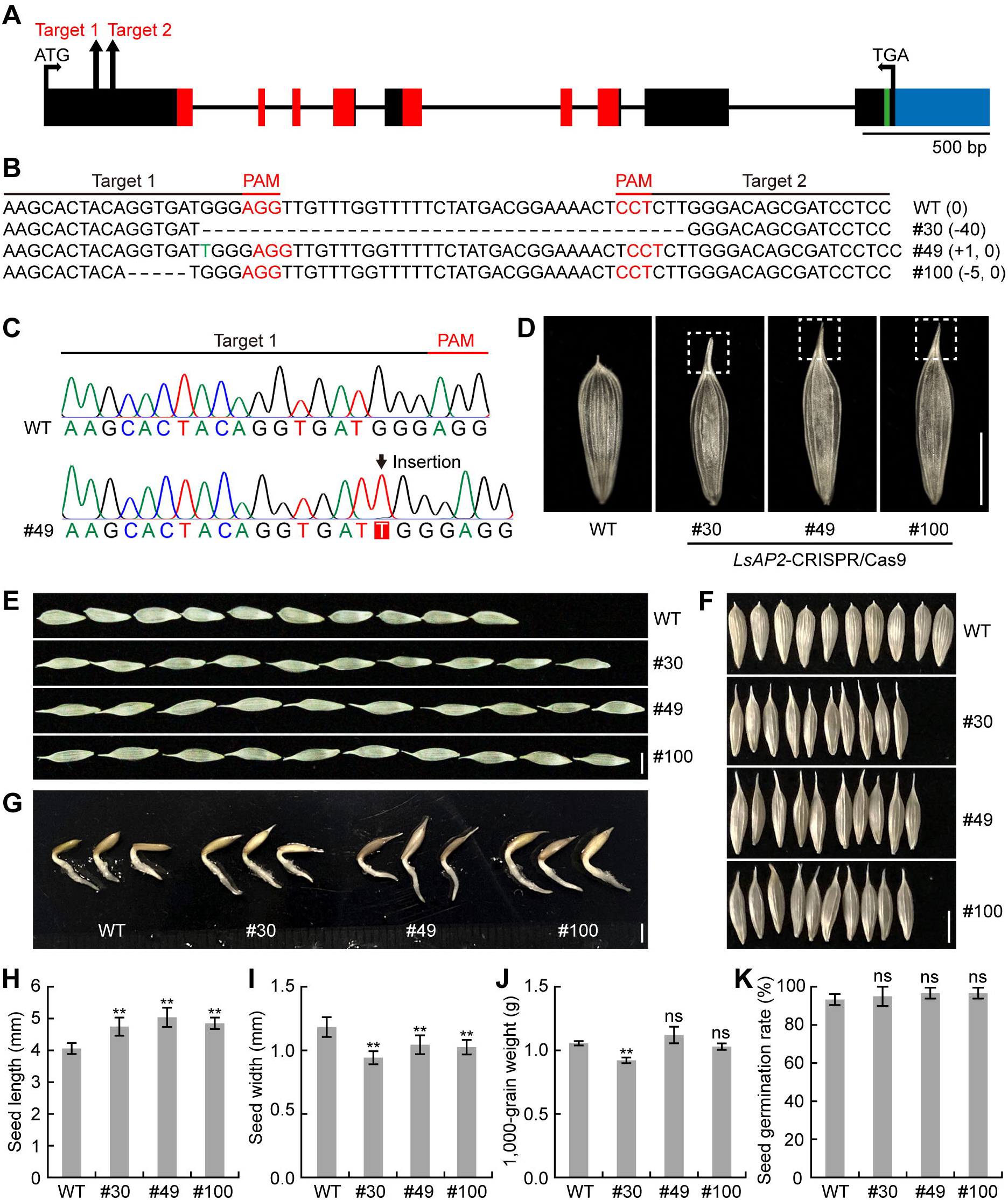
CRISPR/Cas9-mediated editing of *LsAP2* alters seed shape. (A) Gene structure of *LsAP2*. The two target sites are indicated by arrows. Rectangles and lines represent exons and introns, respectively. Red regions indicate the two AP2 domains. Green region indicates the miR172 binding site. Blue region indicates the untranslated region. (B) Sanger sequencing analyses of the mutant alleles in T1 homozygous plants. The target sequences are indicated under the black line. The PAM sequences are depicted in red. The insertion is depicted in green. Numbers on the right side indicate the deletion (−) or insertion (+) compared with WT. (C) Sequencing chromatogram examples of the edited site in the mutant allele compared with WT. Arrow indicates the 1-bp insertion. (D-F) Seed phenotype of WT and three *Lsap2* mutant lines. Extended tips are marked with dotted boxes in D. (G) Seed germination tests of WT and *Lsap2* mutant plants. (H) Seed length of WT and *Lsap2* mutant plants. Values are means ± SD (*n* = 10). (I) Seed width of WT and *Lsap2* mutant plants. Values are means ± SD (*n* = 10). (J) The 1,000-grain weight of WT and *Lsap2* mutant plants. Values are means ± SD (*n* = 5). (K) Seed germination rate of WT and *Lsap2* mutant plants. Values are means ± SD (*n* = 3). Significant difference analysis was conducted with two-tailed Student’s *t* tests (ns, not significant; **, *P* < 0.01). Scale bars: 2 mm.

Given that the CRISPR/Cas9 system was used to edit the genomic sequence of *LsAP2*, genome editing resulted in the non-sense transcripts of *LsAP2* gene. These transcripts disrupted the normal translation of *LsAP2* mRNA with frame-shift mutation and premature stop codons (Supplementary Data S1), which led to the absence of functional LsAP2 protein. *AP2* is an A-class gene that determines perianth identity in flowers (Kunst *et al.*, 1989; Coen and Meyerowitz, 1991). However, we did not observe any defect in the bract, petal, or pappus of *Lsap2* mutant plants. Interestingly, we found that the seed shape was changed in all mutant lines because the seeds were lanky, and there was an extended tip at the top of the seed (Fig. 2D).

In *Lsap2* mutant plants, the seed length increased significantly (Fig. 2E, H), and the seed width decreased significantly (Fig. 2F, I), whereas the 1,000-grain weight did not change significantly compared with WT (Fig. 2J). Seed germination tests showed that almost all seeds germinated, and the roots developed normally after two days incubation at 25°C (Fig. 2G, K), which indicated that the seed germination was not affected in *Lsap2* mutant plants. The scanning electron micrographs revealed that the seed surface of *Lsap2* mutant plants was smoother, and there were fewer longitudinal ribs and striking spines on the seed surface of *Lsap2* mutant plants compared with WT (Supplementary Fig. S5). Further, the length of the seed epidermal cells in *Lsap2* mutant plants was slightly longer than in WT, while the cell width did not change significantly (Supplementary Fig. S5), indicating that the elongated cell length might lead to the increased seed length in *Lsap2* mutant plants.

### Important roles of *LsAP2* in regulating seed morphology

As *LsAP2* was highly expressed at the early stages of seed development, we speculated that LsAP2 might regulate seed shape at the early stages. To verify this idea, we observed the seeds of WT and *Lsap2* mutant plants at different seed development stages. Compared with WT, the seeds of *Lsap2* mutant plants were already longer on the day of pollination (Fig. 3A). Subsequently, the seeds of *Lsap2* mutant plants grew longer and thinner than those of the WT, while the seed beak of *Lsap2* mutant plants was shorter and thicker than that of the WT (Fig. 3B-D). Because the seeds of *Lsap2* mutant plants were lanky, we measured the length-width ratio of the mature seeds and showed that *Lsap2* mutant plants had a larger seed length-width ratio compared with WT (Fig. 3E). Interestingly, in *Lsap2* mutant seeds, the lower part of the beak and upper part of the seed seemed to be fused (Fig. 3B-D). After seed ripening, the beaks were broken and seeds were harvested, and an extended tip was left at the top of the *Lsap2* mutant seeds. These results confirmed that the seed shape of *Lsap2* mutant plants was altered at the early stages of seed development, and this phenotype corresponded to the high expression of *LsAP2* at these stages.

**Fig. 3.**
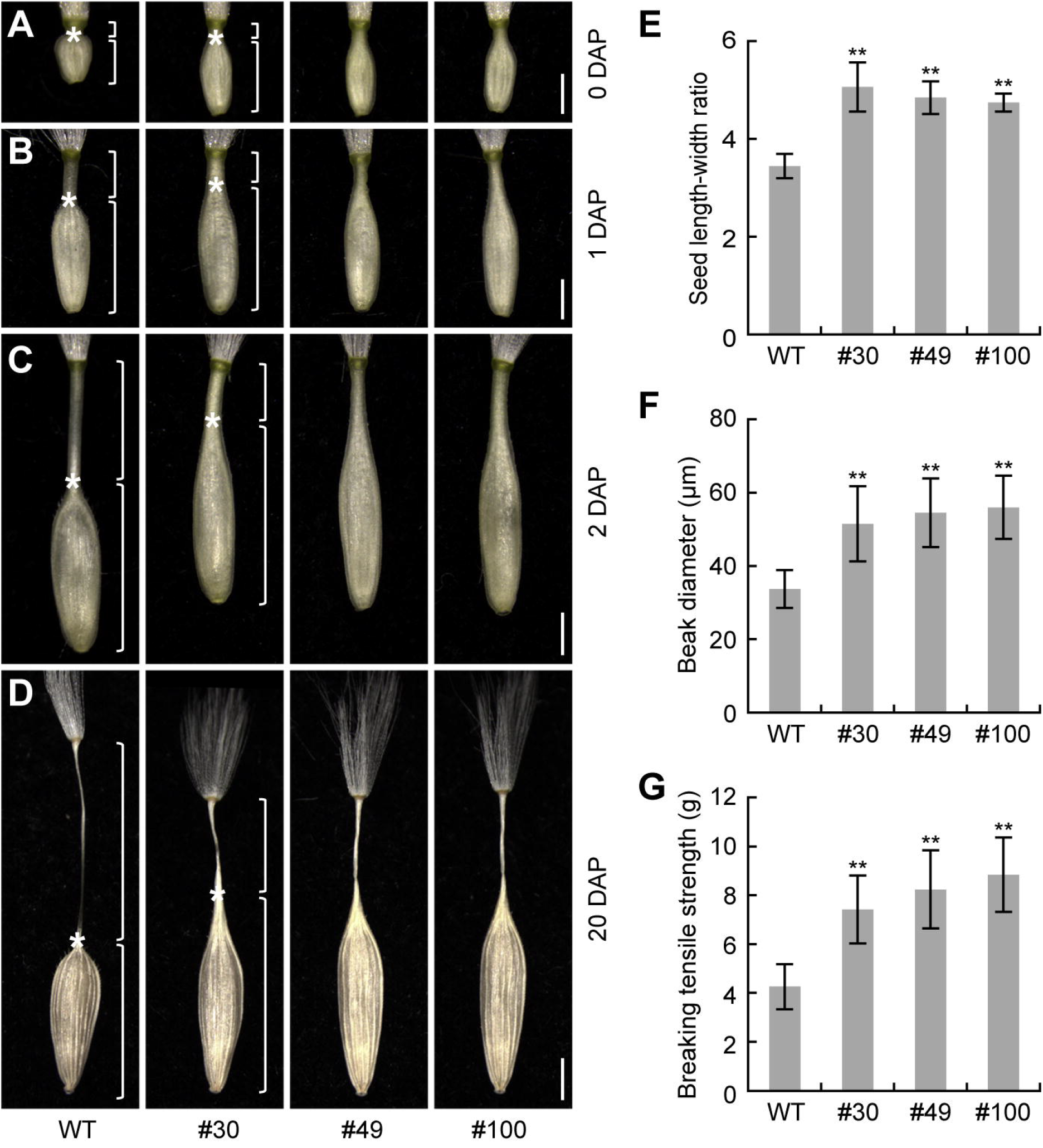
Important roles of *LsAP2* in regulating seed morphology. (A-D) Seed phenotype of WT and three *Lsap2* mutant lines at different seed development stages. Upper and lower brackets indicate beaks and seeds, respectively. Asterisks indicate the junctions between beaks and seeds. (A) The seeds of WT and *Lsap2* mutant plants, 0 DAP. (B) The seeds of WT and *Lsap2* mutant plants, 1 DAP. (C) The seeds of WT and *Lsap2* mutant plants, 2 DAP. (D) The seeds of WT and *Lsap2* mutant plants, 20 DAP. (E) Seed length-width ratio of WT and *Lsap2* mutant plants. Values are means ± SD (*n* = 10). (F) Seed beak diameter of WT and *Lsap2* mutant plants. Values are means ± SD (*n* = 10). (G) The BTS between beak and seed of WT and *Lsap2* mutant plants. Values are means ± SD (*n* = 10). Significant difference analysis was conducted with two-tailed Student’s *t* tests (**, *P* < 0.01). Scale bars: 1 mm.

In *Lsap2* mutant seeds, the beak was thicker and closely connected with the seed, suggesting that this structure might resist stronger external force and contribute to seed dispersal. To verify this hypothesis, we first measured the diameter of the seed beak at the harvest stage, and our data showed that the seed beak diameter of *Lsap2* mutant plants was larger than that of the WT (Fig. 3F). Further, we measured the breaking tensile strength (BTS) between beak and seed, and the results showed that *Lsap2* mutant plants had stronger BTS than WT (Fig. 3G), implying that LsAP2 might have a function in seed dispersal.

### LsAP2-related protein interaction networks during lettuce seed development

Previous studies have shown that AP2 regulates genes related to fruit and seed development. However, whether AP2 could directly interact with these regulators to modulate fruit or seed development are largely unknown. To identify the proteins that potentially interact with LsAP2, we performed yeast two-hybrid assays to test the interactions between LsAP2 and regulators associated with fruit and seed development. Our results showed that LsAP2 directly interacted with four proteins, namely the lettuce homologs of ASYMMETRIC LEAVES2 (AS2), BREVIPEDICELLUS (BP), KANADI1 (KAN1), and KANADI2 (KAN2) (Fig. 4A). *AS2* encodes a LOB domain protein, and AS2 is required for leaf and fruit development in *Arabidopsis* (Iwakawa *et al.*, 2002; Alonso-Cantabrana *et al.*, 2007). *BP* is a class I KNOX gene (Venglat *et al.*, 2002), which modulates replum formation during fruit development in *Arabidopsis* (Alonso-Cantabrana *et al.*, 2007). *KAN1* and *KAN2* are members of the KANADI (KAN) family, and KAN1 and KAN2 have been shown to play important roles in the growth of ovule integuments (McAbee *et al.*, 2006).

**Fig. 4.**
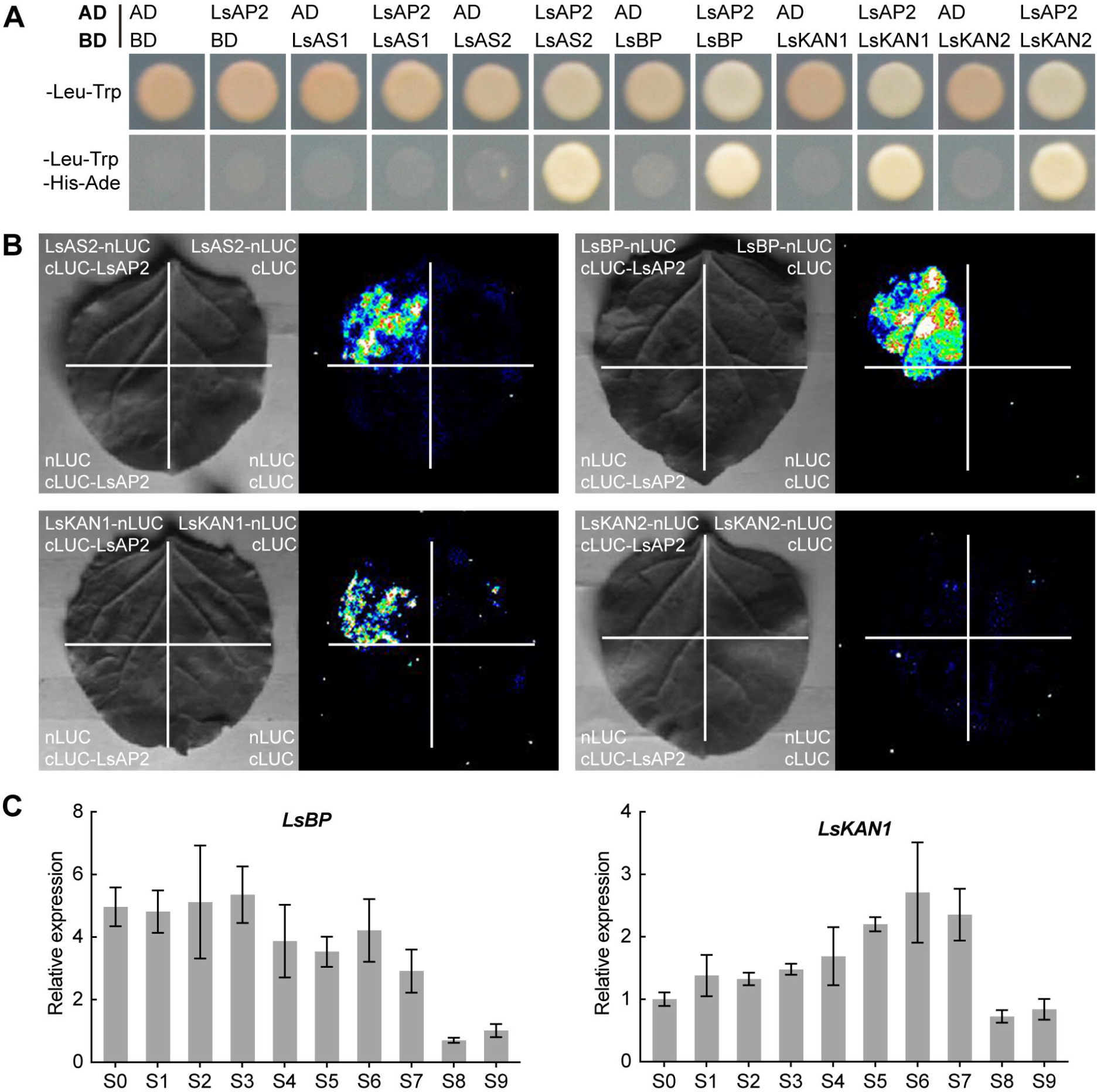
LsAP2-related protein interaction networks during lettuce seed development. (A) LsAP2 interacts with LsAS2, LsBP, LsKAN1, and LsKAN2 in yeast two-hybrid assays. AD, activation domain; BD, binding domain. (B) LCI assays show that LsAP2 interacts with LsAS2, LsBP, and LsKAN1 *in planta*. (C) Expression of *LsBP* and *LsKAN1* at different seed development stages. Values are means ± SD (*n* = 3). The expression data were normalized using *LsPP2A-1* and *LsTIP41* as reference genes.

To confirm these potential interactions *in planta*, we performed LCI assays in tobacco leaves. Co-expression of LsAP2 with LsAS2, LsBP, or LsKAN1 produced strong fluorescence activity (Fig. 4B), whereas co-expression of LsAP2 with LsKAN2 or control vectors showed only background levels of fluorescence activity (Fig. 4B). These results demonstrated that LsAP2 could interact with LsAS2, LsBP, and LsKAN1 *in planta*.

To determine whether the interactions of LsAP2 with LsAS2, LsBP, and LsKAN1 might be important for seed development, we performed expression analyses of *LsAS2*, *LsBP*, and *LsKAN1*. Our data showed that *LsBP* and *LsKAN1* were expressed in seeds, while *LsAS2* was not detected in seeds. Significantly, *LsBP* was strongly expressed in seeds (Supplementary Fig. S6). Further experiments were performed to examine the expression levels of *LsBP* and *LsKAN1* at different seed development stages. The qRT-PCR analysis showed that *LsBP* was expressed at all stages and displayed higher expression level at the early stages, and the expression of *LsBP* decreased with the development of the seeds (Fig. 4C), which was overlapped temporally with the expression pattern of *LsAP2*. The transcripts of *LsKAN1* could also be detected at different developmental stages, but *LsKAN1* exhibited a different expression pattern from *LsAP2* (Fig. 4C). The biological relevance between *LsAP2* and *LsBP* indicated that LsAP2 might cooperate with LsBP to regulate seed shape.

### LsAP2 affects the expression of fruit development related genes

During the fruit development of *Arabidopsis*, AP2 negatively regulates *BP* and *REPLUMLESS* (*RPL*) to prevent replum overgrowth. Meanwhile, AP2 negatively regulates valve margin formation by repressing the expression of valve margin identity genes (Ripoll *et al.*, 2011). Since the seeds of lettuce are also fruits, and the morphology of *Lsap2* mutant seeds was changed, we then examined the expression of fruit development related genes in *Lsap2* mutant plants. Compared with WT, the expression levels of *LsBP* and *LsRPL* were significantly decreased in the seeds of *Lsap2* mutant plants (Fig. 5A). However, orthologs of *SHATTERPROOF1* (*SHP1*), *SHATTERPROOF2* (*SHP2*), *INDEHISCENT* (*IND*), and *ALCATRAZ* (*ALC*), which are required for valve margin formation and dehiscence, had not been found in lettuce. The phylogenetic analysis revealed that LsAP2 was closely related to *Arabidopsis* AP2, and we demonstrated that LsAP2 also functioned in lettuce seed and fruit development. However, our results implied that LsAP2 might positively regulate *LsBP* and *LsRPL*, which was opposite from *Arabidopsis*. Actually, the fruit structure of lettuce is different from that of *Arabidopsis*. The achenes of lettuce are indehiscent fruits, whereas the siliques of *Arabidopsis* are dehiscent fruits. Similarly, the caryopses of rice are also dry fruits with indehiscent, and the AP2-like transcription factor SNB positively regulates the expression of two rice *RPL* orthologs, *qSH1* and *SH5* (Jiang *et al.*, 2019). These results indicated that the *AP2* genes might exhibit divergent regulatory patterns between different types of fruit because of the structural differences.

**Fig. 5.**
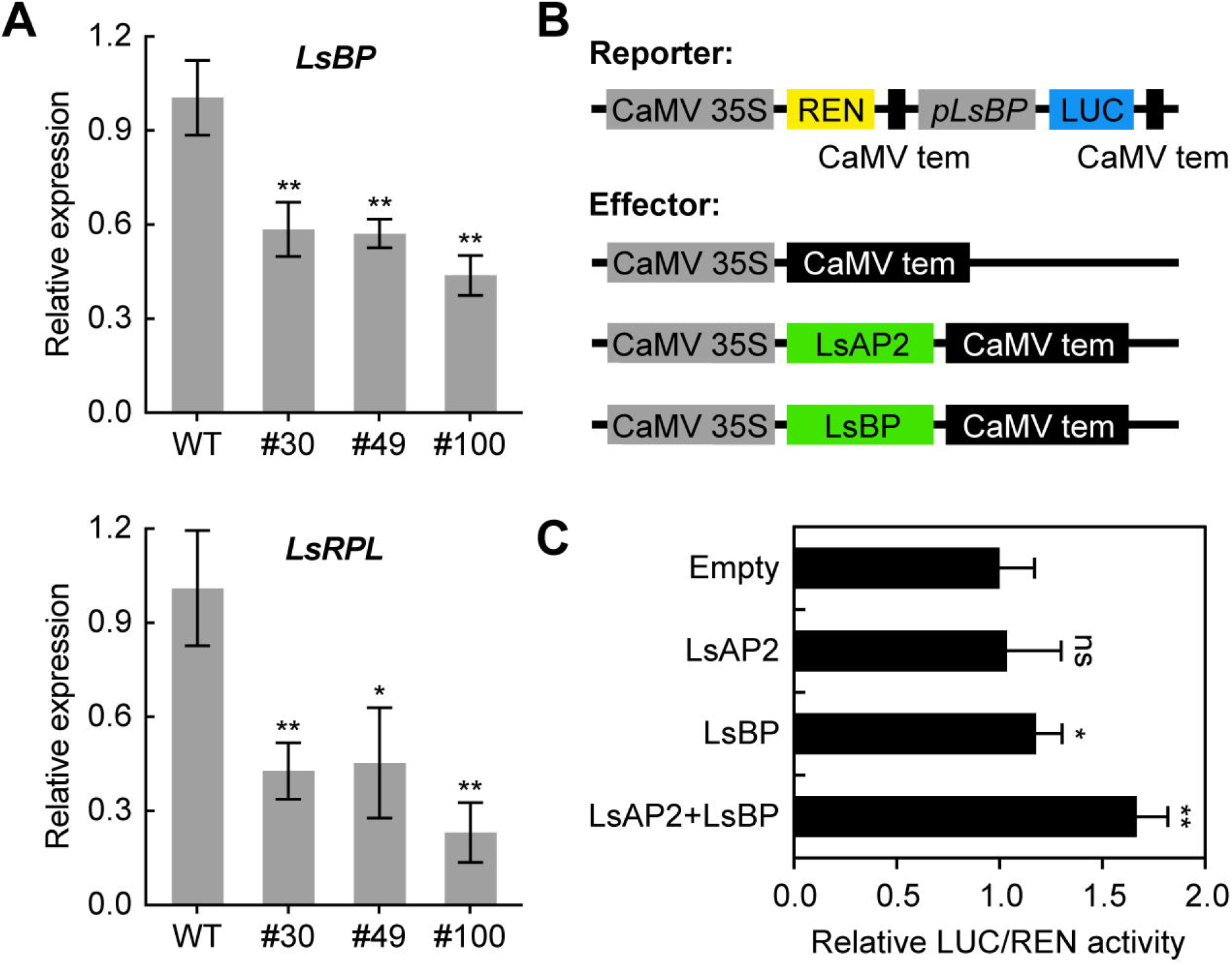
LsAP2 affects the expression of fruit development related genes. (A) Expression of *LsBP* and *LsRPL* between WT and *Lsap2* mutant plants. Values are means ± SD (*n* = 3). The expression data were normalized using *LsPP2A-1* and *LsTIP41* as reference genes. (B) Schematic diagram of the reporter and effector constructs. (C) Dual-luciferase reporter assays show the interactions of LsAP2 and LsBP with the *LsBP* promoter. Values are means ± SD (*n* = 6). Significant difference analysis was conducted with two-tailed Student’s *t* tests (ns, not significant; *, *P* < 0.05; **, *P* < 0.01).

Interestingly, LsAP2 interacted with LsBP at the protein level, and the *LsBP* gene was also down-regulated in *Lsap2* mutant plants. We speculated that LsBP or LsAP2 might promote *LsBP* expression, and the protein interaction between LsAP2 and LsBP would enhance the promotion. To test this regulatory network, we used the *LsBP* promoter to drive *LUC* reporter gene and performed dual-luciferase reporter assays. Compared with the control experiment, expression of LsAP2 or LsBP alone had a minor effect on the promotion of LUC activity, while co-expression of LsAP2 and LsBP significantly increased the LUC activity (Fig. 5C), indicating that LsAP2 cooperated with LsBP to enhance the activity of *LsBP* promoter. These results demonstrated that the protein interaction between LsAP2 and LsBP could promote the expression of *LsBP*, which might be important for seed development in lettuce.

### Identification of differentially expressed genes between WT and *Lsap2* mutant plants

To dissect the molecular function of *LsAP2* in seed development further, we performed the transcriptome analysis of the seeds from WT and *Lsap2* mutant plants on the day of pollination. Many genes were differentially expressed between WT and *Lsap2* mutant plants, and we identified 586 up-regulated genes and 547 down-regulated genes among the transcriptomes (Supplementary Table S6). GO enrichment analysis of the DEGs revealed that they were involved in multiple processes. Significantly enriched GO terms included nucleosome assembly and its related cellular component. Moreover, some GO categories were related to photosynthesis (Fig. 6A). Nucleosome assembly is involved in DNA replication and cell division, and seed photosynthesis is essential for storage activity. These processes are necessary for normal seed development.

**Fig. 6.**
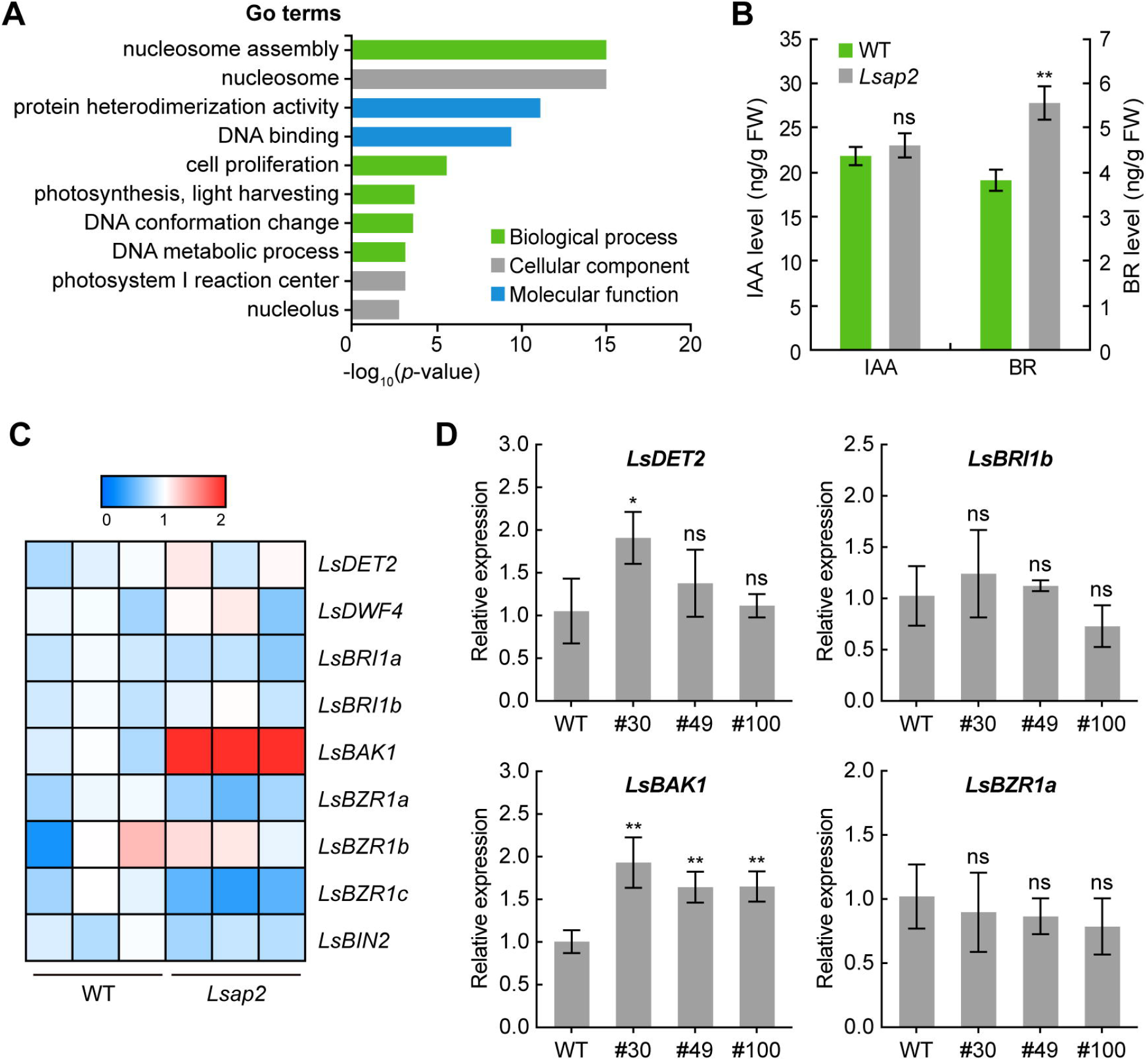
Identification of differentially expressed genes between WT and *Lsap2* mutant plants. (A) GO enrichment analysis of the DEGs. (B) The IAA and BR levels in the developing seeds of WT and *Lsap2* mutant plants. Values are means ± SD (*n* = 3). (C) Heat map for the expression levels of BR-related genes between WT and *Lsap2* mutant plants. Red boxes indicate up-regulation, and blue boxes indicate down-regulation. (D) Expression analyses of BR-related genes between WT and *Lsap2* mutant plants. Values are means ± SD (*n* = 3). The expression data were normalized using *LsPP2A-1* and *LsTIP41* as reference genes. Significant difference analysis was conducted with two-tailed Student’s *t* tests (ns, not significant; *, *P* < 0.05; **, *P* < 0.01).

Further KEGG analysis of the DEGs indicated that some hormone-related genes were differentially expressed between WT and *Lsap2* mutant plants, such as cytokinin, auxin, and BR. Phytohormones have various functions in plant growth and development. Because auxin and BR have been shown to regulate seed growth (Jiang *et al.*, 2013; Figueiredo and Köhler, 2018), we measured the endogenous hormones in the developing seeds of WT and *Lsap2* mutant plants. We found that the BR level was significantly increased in the developing seeds of *Lsap2* mutant plants compared with WT. However, there was no significant difference in IAA level between WT and *Lsap2* mutant plants (Fig. 6B).

To explore the significance of the increased BR level in *Lsap2* mutant seeds, we examined the expression of BR-related genes that have been shown to control seed development. The transcriptome data indicated that a lettuce homolog of *BRASSINOSTEROID INSENSITIVE 1–ASSOCIATED KINASE 1* (*BAK1*) was significantly up-regulated in *Lsap2* mutant plants (Fig. 6C), while the expression levels of other BR-related genes were not changed significantly. We also performed qRT-PCR assays to verify the transcription levels of some genes, and the results were generally consistent with the transcriptome data (Fig. 6D). BAK1 is a receptor kinase involved in BR signal transduction (Li *et al.*, 2002), and it has been shown to regulate grain size in rice. Loss of function of *OsBAK1* exhibits decreased grain size (Yuan *et al.*, 2017). Thus, we speculated that the increased BR level in *Lsap2* mutant seeds might stimulate the BR perception and signaling, thus promoting cell elongation and increasing seed length.

## Discussion

### Unique roles of *LsAP2* during seed development in lettuce

*AP2* gene encodes a transcription factor, and the broad roles of the *AP2* gene in plant growth and development have been shown in many species. Here, we identified a lettuce *AP2* gene and explored its biological function. Expression analysis revealed that *LsAP2* was highly expressed in seeds, especially at the early stages of seed development (Fig. 1C). We used the CRISPR/Cas9 system to edit *LsAP2* gene and found that LsAP2 regulates lettuce seed shape (Fig. 2A-D). Although the loss of *AP2* function alters seed shape in both *Arabidopsis* and lettuce, there are some differences. In *Arabidopsis*, *ap2* mutants produce large seeds with aberrant shapes. Reducing *AP2* gene activity increases seed length and width and leads to increased seed mass (Jofuku *et al.*, 2005; Ohto *et al.*, 2005). Meanwhile, the epidermal cells of *ap2* mutant seeds lack the columella structure, and the epidermal cells are larger and more irregular in shape than WT seeds (Jofuku *et al.*, 1994). In lettuce, the seed shape of *Lsap2* mutants was affected as the seeds were lanky and developed an extended tip (Fig. 2D-F). In *Lsap2* mutant plants, the seed length increased, and the seed width decreased (Fig. 2H, I), while the seed mass did not change significantly compared with WT (Fig. 2J). Our scanning electron micrographs showed that the seed surface of *Lsap2* mutants was smoother than that of the WT due to decreased longitudinal ribs and fewer striking spines. Moreover, cell length of the seed epidermal cells increased slightly in *Lsap2* mutant plants, while the cell shape and cell structure did not show noticeable changes (Supplementary Fig. S5).

In the Asteraceae plant dandelion, the pappus attaches to the beak for seed dispersal, and the seed falls and stops dispersing when the junction between beak and seed is broken. Lettuce also belongs to the Asteraceae family and has the same seed structure as dandelion. In *Lsap2* mutant seeds, the beak was thicker than that of the WT (Fig. 3A-D). Besides, the beak was closely connected with the seed top, and the BTS between beak and seed was stronger than WT (Fig. 3G), implying that this structure may resist stronger external force and contribute to longer distance seed dispersal with the help of the wind. This finding may also reveal a potential new function of *LsAP2* in seed dispersal, which is radically divergent from *Arabidopsis* as the seed structure of lettuce is different from *Arabidopsis*.

Since *AP2* is an A-class gene that determines the identity of perianth organs, the sepals and petals transform into reproductive organs in *ap2* mutants (Kunst *et al.*, 1989; Coen and Meyerowitz, 1991). Although the expression analysis showed that *LsAP2* was expressed in bract and petal (Fig. 1B), we did not observe any perianth defect or transformation in the floral organs of *Lsap2* mutant plants. On one hand, the petunia AP2-type *REPRESSOR OF B-FUNCTION* (*ROB*) genes have incomplete A-function. The *ROB* genes are redundantly required for the normal development of the perianth and are not required to antagonize the C-function in the perianth (Morel *et al.*, 2017). Based on current understanding, it is also possible that the complete A-function of *AP2* gene does not occur in lettuce. On the other hand, the lettuce reference genome was published in 2017, and researchers found that some types of genes, including *AP2*, were enriched in the triplicated regions (Reyes-Chin-Wo *et al.*, 2017). Indeed, there are three *AP2* genes in lettuce (Supplementary Fig. S1), and this gene redundancy mechanism may explain the absence of perianth defect in *Lsap2* mutant plants.

### LsAP2 may regulate seed shape via interacting with LsBP

In *Arabidopsis*, the regulatory networks of fruit development have been studied extensively, and many genetic regulators involved in fruit development were identified. BP and RPL play direct roles in promoting replum identity during fruit development. AP2 cooperates with AS1 and AS2 to modulate replum development by independently regulating the activity of *BP* and *RPL* (Ripoll *et al.*, 2011). Although their interactions are known at the genetic level, the direct protein interaction networks of AP2 in fruit or seed development are largely unknown. In this study, we identified several interactors of LsAP2 using yeast two-hybrid and LCI assays. LsAP2 directly interacted with LsBP *in vivo* (Fig. 4A, B), and expression analysis revealed that *LsBP* was highly expressed in seeds and showed a temporally overlapping expression pattern with *LsAP2* (Fig. 4C; Supplementary Fig. S6).

As LsAP2 directly interacted with LsBP at the protein level, and the expression of *LsBP* gene was also decreased in *Lsap2* mutant seeds (Fig. 5A), we further performed dual-luciferase reporter assays to explore the regulatory network between LsAP2 and LsBP. Our results indirectly demonstrated that the protein interaction between LsAP2 and LsBP increased the gene expression of *LsBP* (Fig. 5C). Together, the high biological relevance between *LsAP2* and *LsBP* suggests that LsAP2 may cooperate with LsBP to regulate seed shape in lettuce.

### LsAP2 may negatively regulate BR biosynthesis and signaling in seeds

Previous studies showed that BR plays crucial roles in regulating seed size and seed shape (Jiang *et al.*, 2013; Song, 2017). BR functions through the cell surface receptor BRASSINOSTEROID INSENSITIVE1 (BRI1) and the transcription factor BRASSINAZOLE-RESISTANT1 (BZR1) to control BR-responsive genes (Nolan *et al.*, 2020). In both *Arabidopsis* and rice, inactivation of BR biosynthesis genes or BR receptors lead to reduced seed size and abnormal seed shape, while elevated BR content or enhanced BR signaling results in increased seed size. In *Arabidopsis*, the seeds of the BR-deficient mutant *de-etiolated2* (*det2*) and the BR-insensitive mutant *bri1-5* are smaller and less elongated than those of the WT (Jiang *et al.*, 2013). A rice gain-of-function mutant, *slender grain Dominant* (*slg-D*), shows increased BR level and longer grains (Feng *et al.*, 2016), while the loss of function of *OsBAK1* (Yuan *et al.*, 2017), a receptor kinase involved in BR signaling, results in reduced grain length. In this study, we found that the BR content of the *Lsap2* mutant seeds was much higher than that of the WT (Fig. 6B), and the transcription level of *LsBAK1* increased significantly in *Lsap2* mutant seeds (Fig. 6C, D). Meanwhile, the seed length and seed length-width ratio of *Lsap2* mutant plants were larger than those of the WT (Fig. 2H; Fig. 3E). Our results are consistent with the reports that elevated BR content or enhanced BR signaling results in increased seed length. Thus, we speculate that LsAP2 may negatively regulate BR biosynthesis and signaling to modulate seed morphology.

Taken together, LsAP2 has a significant function during seed development in lettuce, and we proposed a hypothetical working model in which LsAP2 may regulate lettuce seed shape via interacting with LsBP and repressing BR biosynthesis and signaling (Fig. 7). Applying the CRISPR/Cas9 system to generate the knockout plants of the identified interactors or downstream genes of LsAP2 will shed light on the unique regulatory networks during lettuce seed development.

**Fig. 7.**
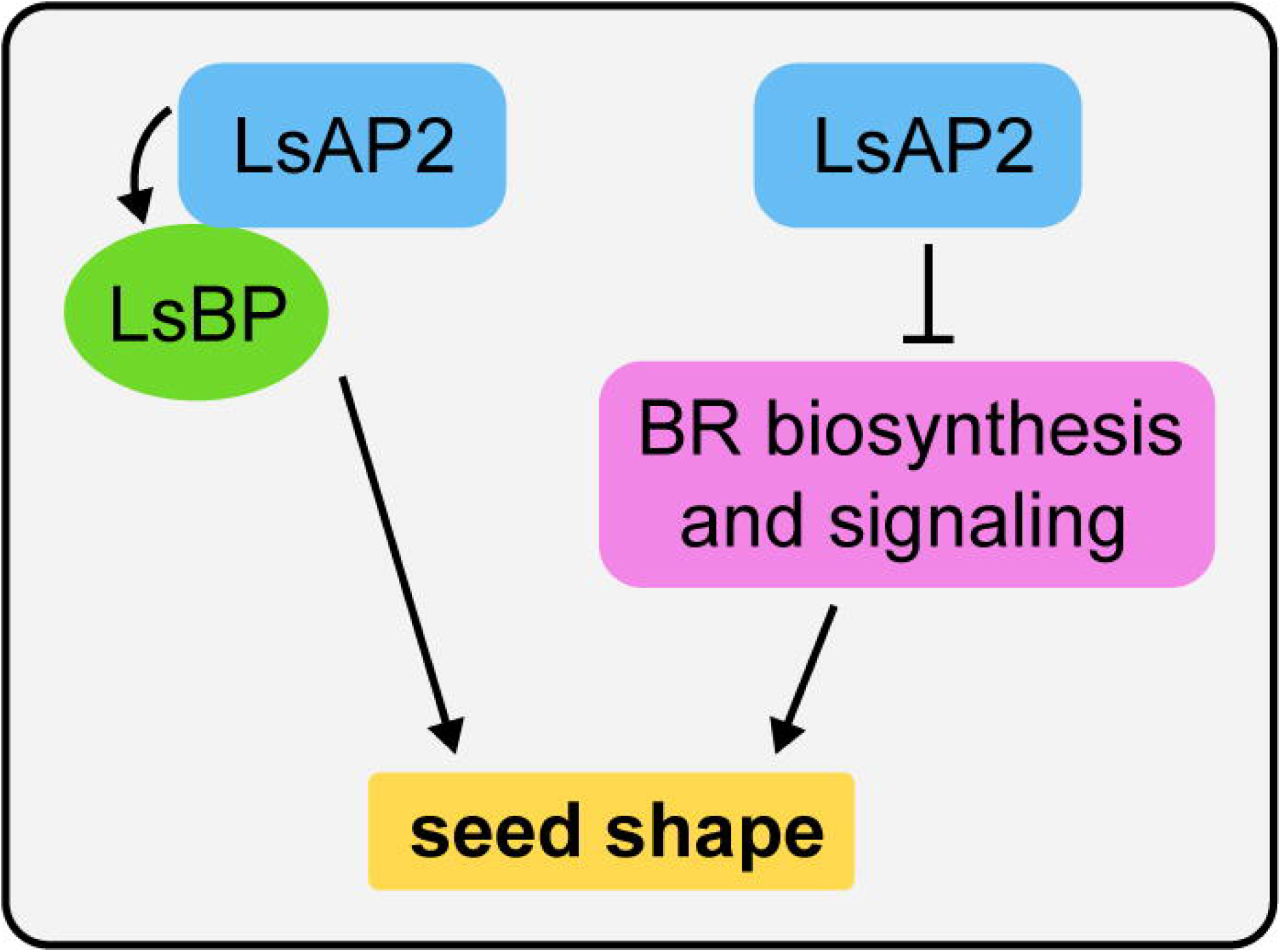
A hypothetical working model of LsAP2 in regulating seed shape in lettuce. LsAP2 may regulate seed shape by directly interacting with LsBP and promoting the expression of *LsBP*. Meanwhile, LsAP2 may negatively regulate BR biosynthesis and signaling in lettuce seeds. The biological function of *LsBP* needs to be further confirmed in lettuce. Arrow, positive regulation; bar, negative regulation.

## Supplementary data

Supplementary data are available at *JXB* online.

Fig. S1. Phylogenetic tree of the euAP2 subfamily.

Fig. S2. Expression of *LsAP2* in different tissues.

Fig. S3. Sanger sequencing analyses of the mutant alleles in T0 generation plants.

Fig. S4. Sequencing chromatogram analyses of the edited sites in T0 generation plants.

Fig. S5. Scanning electron microscope observation of the seeds from WT and *Lsap2* mutant plants.

Fig. S6. Expression analyses of the interactors of LsAP2. Table S1. Gene information in this study.

Table S2. Primers used in this study.

Table S3. FPKM values of lettuce *AP2* genes at different seed development stages. Table S4. Editing efficiency and mutation types of CRISPR/Cas9-mediated editing of *LsAP2* in T0 generation plants.

Table S5. Summary of the putative off-target sites analyses. Table S6. List of DEGs between WT and *Lsap2* mutant plants.

Data S1. Coding sequences and amino acid sequences of *LsAP2* gene in WT and three mutant lines.

## Acknowledgments

We thank Dr. Shuangxi Fan and Dr. Yingyan Han (Beijing University of Agriculture) for providing the experimental materials. We are grateful to Dr. Xiaolan Zhang (China Agricultural University) for technical assistance. This work was supported by Beijing Leafy Vegetables Innovation Team of Modern Agro-industry Technology Research System (BAIC07-2020) and The Construction of Beijing Science and Technology Innovation and Service Capacity in Top Subjects (CEFF-PXM2019_014207_000032).

## Author contributions

QW and CL designed the project. CL, SW, KN, ZC, YW, JY and MQ performed the experiments. CL, SW and KN analyzed the experimental data. QW and CL wrote the manuscript with the help of SW, KN and ZC. All the authors read and approved the final manuscript.

## Data availability statement

All data supporting the findings of this study are available within the paper and within its supplementary data published online.

## Notes

### Competing Interest Statement

The authors have declared no competing interest.

### Summary of Updates

We have addressed the outstanding points and carefully revised our manuscript. The quality of the written English was also improved. We repeated the qRT-PCR assays using LsPP2A-1 and LsTIP41 as reference genes to normalize the expression data. The new results were generally consistent with our previous results. The new data were presented in our revised figures (Fig. 1B, C; Fig. 4C; Fig. 5A; Fig. 6D; Supplementary Figs. S2, S6), and the method was modified in the 'Materials and methods' section of the revised manuscript. We also stated the number of biological replicates where mean and SD are given.

