## Supplementary Figures S1-S6+Tables S1-S5+Data S1 for "The APETALA2 transcription factor LsAP2 regulates seed shape in lettuce"

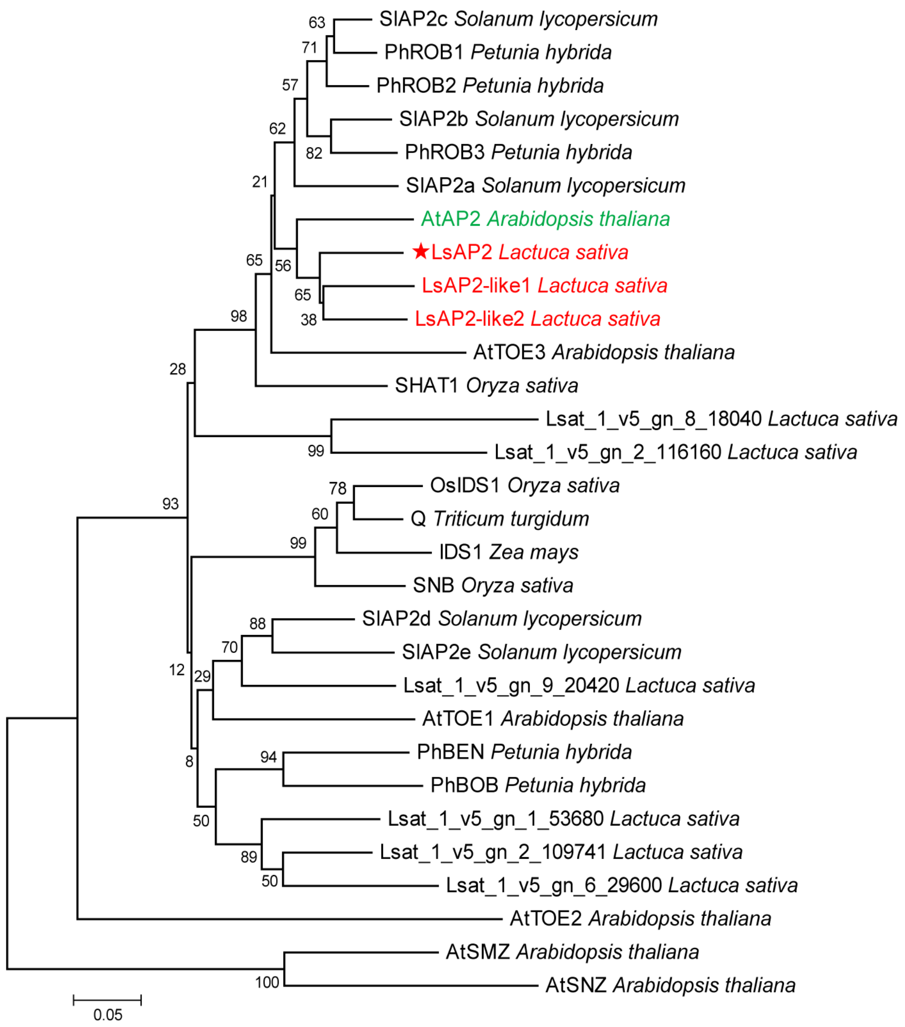


**Fig. S1. Phylogenetic tree of the euAP2 subfamily.** The amino acid sequences are from *Arabidopsis* (At), lettuce (Ls), tomato (Sl), petunia (Ph), rice (Os), maize (Zm), and wheat (Tt). *Arabidopsis* AP2 is depicted in green font, lettuce *AP2* genes are depicted in red font, and an asterisk is in front of the LsAP2. Bootstrap values (%) based on 1000 replicates are indicated near the branching points.

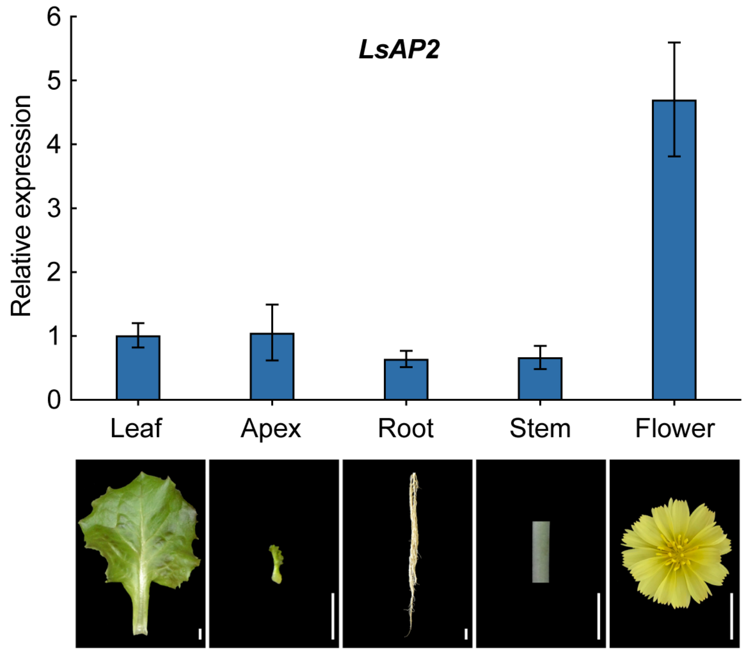


**Fig. S2. Expression of *LsAP2* in different tissues.** Images under the histogram correspond to the different lettuce tissues. Values are means ± SD ($n$ = 3). The expression data were normalized using *LsPP2A-1* and *LsTIP41* as reference genes. Scale bars: 1 cm.

```
                 Target 1         PAM                                   PAM         Target 2
WT       AAGCACTACAGGTGAT-GGGAGGTTGTTTGGTTTTTCTATGACGGAAAACTCCTCTTGGGACAGCGATCCTCC  WT
#24-A1   AAGCACTACAGGTGA--GGGAGGTTGTTTGGTTTTTCTATGACGGAAAACTCCTCTTGGGACAGCGATCCTCC  -1, 0
   -A2   AAGCACTACAGGT--T-GGGAGGTTGTTTGGTTTTTCTATGACGGAAAACTCCTCTTGGGACAGCGATCCTCC  -2, 0
#30-A1   AAGCACTACAGGTGAT-----------------------------------------GGGACAGCGATCCTCC  -40
   -A2   AAGCACTACAGGT----GGGAGGTTGTTTGGTTTTTCTATGACGGAAAACTCCTCTT--GACAGCGATCCTCC  -3, -2
#49      AAGCACTACAGGTGATTGGGAGGTTGTTTGGTTTTTCTATGACGGAAAACTCCTCTTGGGACAGCGATCCTCC  +1, 0
#59      AAGCACTACAGGTGATTGGGAGGTTGTTTGGTTTTTCTATGACGGAAAACTCCTCTTGGGACAGCGATCCTCC  +1, 0
#100-A1  AAGCACTACA-----T-GGGAGGTTGTTTGGTTTTTCTATGACGGAAAACTCCTCTTGGGACAGCGATCCTCC  -5, 0
    -A2  AAGCACTACAGGTGATTGGGAGGTTGTTTGGTTTTTCTATGACGGAAAACTCCTCTTGGGACAGCGATCCTCC  +1, 0
#165-A1  AAGCACTACAGG-----GGGAGGTTGTTTGGTTTTTCTATGACGGAAAACTCCTCTTGGGACAGCGATCCTCC  -4, 0
    -A2  AAGCACTACAGGTGATTGGGAGGTTGTTTGGTTTTTCTATGACGGAAAACTCCTCTTGGGACAGCGATCCTCC  +1, 0
```

**Fig. S3. Sanger sequencing analyses of the mutant alleles in T0 generation plants.** The target sequences are indicated under the black line. The PAM sequences are depicted in red. The insertions are depicted in green. Numbers on the right side indicate the deletion (-) or insertion (+) compared with WT.

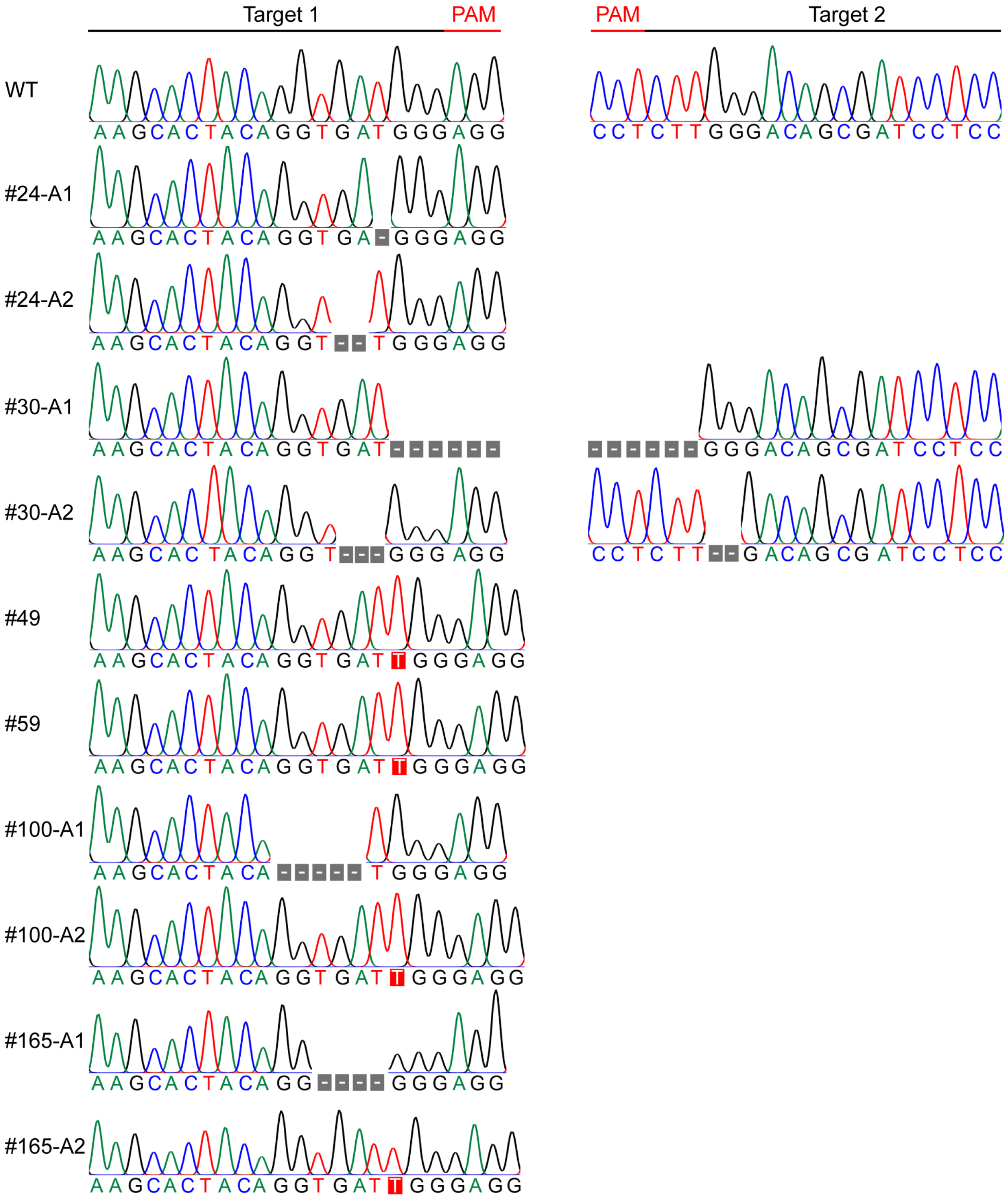


**Fig. S4. Sequencing chromatogram analyses of the edited sites in T0 generation plants.**

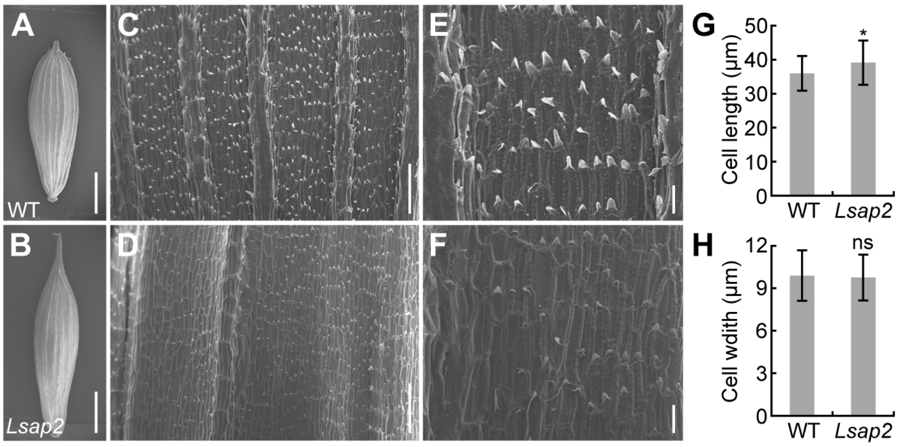


**Fig. S5. Scanning electron microscope observation of the seeds from WT and *Lsap2* mutant plants.** (A, B) SEM images of the mature seeds from WT and *Lsap2* mutant plants. (C, D) SEM images of the seed epidermal cells from WT and *Lsap2* mutant plants. (E, F) Magnified images of the seed epidermal cells from C, D, respectively. (G) Cell length of seed epidermal cells from WT and *Lsap2* mutant plants. Values are means ± SD ($n$ = 30). (H) Cell width of seed epidermal cells from WT and *Lsap2* mutant plants. Values are means ± SD ($n$ = 30). Significant difference analysis was conducted with two-tailed Student's $t$ tests (ns, not significant; *, $P < 0.05$). Scale bars: (A, B) 1 mm; (C, D) 100 µm; (E, F) 20 µm.

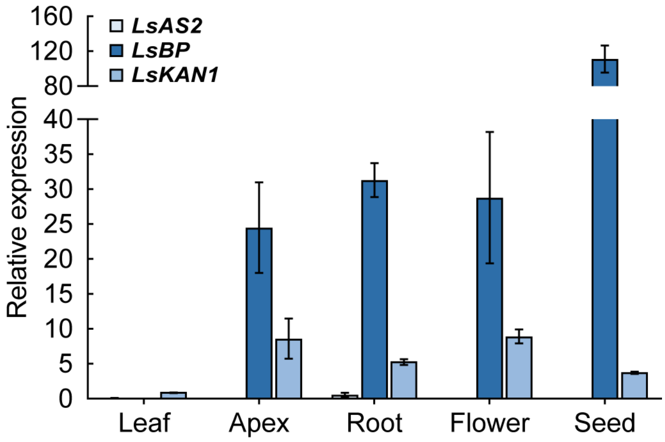


**Fig. S6. Expression analyses of the interactors of LsAP2.** Values are means ± SD ($n$ = 3). The expression data were normalized using *LsPP2A-1* and *LsTIP41* as reference genes.

**Table S1. Gene information in this study.**

| Species | Gene name | Accession number |
|---|---|---|
| *Lactuca sativa* | *LsAP2* | Lsat_1_v5_gn_3_89681 |
| | *LsAP2-like1* | Lsat_1_v5_gn_4_165341 |
| | *LsAP2-like2* | Lsat_1_v5_gn_2_85920 |
| | *LsPP2A-1* | Lsat_1_v5_gn_8_160621 |
| | *LsTIP41* | Lsat_1_v5_gn_5_116421 |
| | *LsAS1* | Lsat_1_v5_gn_4_153581 |
| | *LsAS2* | Lsat_1_v5_gn_2_108440 |
| | *LsBP* | Lsat_1_v5_gn_6_108340 |
| | *LsKAN1* | Lsat_1_v5_gn_3_67301 |
| | *LsKAN2* | Lsat_1_v5_gn_3_18021 |
| | *LsRPL* | Lsat_1_v5_gn_9_33021 |
| | *LsDET2* | Lsat_1_v5_gn_5_155401 |
| | *LsDWF4* | Lsat_1_v5_gn_7_92481 |
| | *LsBRI1a* | Lsat_1_v5_gn_4_138061 |
| | *LsBRI1b* | Lsat_1_v5_gn_2_67280 |
| | *LsBAK1* | Lsat_1_v5_gn_9_117520 |
| | *LsBZR1a* | Lsat_1_v5_gn_2_79760 |
| | *LsBZR1b* | Lsat_1_v5_gn_4_122280 |
| | *LsBZR1c* | Lsat_1_v5_gn_5_29981 |
| | *LsBIN2* | Lsat_1_v5_gn_1_109401 |
| *Arabidopsis thaliana* | *AtAP2* | AT4G36920 |
| | *AtTOE3* | AT5G67180 |
| | *AtTOE1* | AT2G28550 |
| | *AtTOE2* | AT5G60120 |
| | *AtSMZ* | AT3G54990 |
| | *AtSNZ* | AT2G39250 |
| *Solanum lycopersicum* | *SlAP2a* | Solyc03g044300 |
| | *SlAP2b* | Solyc02g064960 |
| | *SlAP2c* | Solyc02g093150 |
| | *SlAP2d* | Solyc11g072600 |
| | *SlAP2e* | Solyc06g075510 |
| *Petunia hybrida* | *PhROB1* | Peaxi162Scf00549g00114 |
| | *PhROB2* | Peaxi162Scf00685g00012 |
| | *PhROB3* | Peaxi162Scf02748g00006 |
| | *PhBEN* | Peaxi162Scf00005g00506 |
| | *PhBOB* | Peaxi162Scf00472g00069 |
| *Zea mays* | *IDS1* | AF048900 |
| *Oryza sativa* | *OsIDS1* | LOC_Os03g60430 |
| | *SNB* | LOC_Os07g13170 |
| | *SHAT1* | LOC_Os04g55560 |

| | | |
|---|---|---|
| *Triticum turgidum* | *Q* | AY702960 |

**Table S2. Primers used in this study.**

| Purpose | Primer sequence (5′−3′) |
|---|---|
| **qRT-PCR** | |
| LsAP2-qRT-F | GTGGTGGCTCGGCTTCAAATA |
| LsAP2-qRT-R | GGTCCATAAAATGCGATGTCCTT |
| LsAS2-qRT-F | AATGTTCACAAGGTGTTCGGA |
| LsAS2-qRT-R | TGGTGTTGGAGCAGGGAGAT |
| LsBP-qRT-F | CAACAAAAGGACAACAATTCGC |
| LsBP-qRT-R | GGGAGAGAAAGAAAGGATCGAT |
| LsKAN1-qRT-F | CAATTTGTCCTTGTAACGGACA |
| LsKAN1-qRT-R | CACAATCCAAAGAAGTCAAGGG |
| LsRPL-qRT-F | TTGAACTTGCACAGTTATGGTG |
| LsRPL-qRT-R | ACCTTTCAAAATCGAAGCGTAG |
| LsDET2-qRT-F | TCTCAAATGGGTATCCGATCAG |
| LsDET2-qRT-R | CTCTCGGAATCTTATACCCACC |
| LsBRI1b-qRT-F | AATCCGTCGAGAGGATTTAGAC |
| LsBRI1b-qRT-R | TCGGACCTGTAAGATCGTTATG |
| LsBAK1-qRT-F | GTCATCAAGACAATCACGAAGG |
| LsBAK1-qRT-R | CATTGAACACGGCTTAGTCAAA |
| LsBZR1a-qRT-F | CACAACCGCATTACAGTCATTC |
| LsBZR1a-qRT-R | TTTTCACAAGGTTGAAAGTCGG |
| LsPP2A-1-qRT-F | ATTCATGGTCAATTCTACGATCTGGT |
| LsPP2A-1-qRT-R | GAATAATACCCGCGATCAACATAATC |
| LsTIP41-qRT-F | TTTGTATGGAGATGAATTGGCTGATA |
| LsTIP41-qRT-R | CGTAAGAGAAGAAACCAACAGCTAGG |
| **Lettuce transformation** | |
| pLsAP2-GUS-pBI121-F | GATTACGCCAAGCTTAGTACGTTGCTATTAATAACC |
| pLsAP2-GUS-pBI121-R | ACCACCCGGGGATCCTTCTCCCTCCAAAACCAACA |
| LsAP2-CRISPR/Cas9-BsF | ATATATGGTCTCGATTGAAGCACTACAGGTGATGGGGTT |
| LsAP2-CRISPR/Cas9-BsR | ATTATTGGTCTCGAAACCTTGGGACAGCGATCCTCCCAA |
| LsAP2-CRISPR/Cas9-F0 | TGAAGCACTACAGGTGATGGGGTTTTAGAGCTAGAAATAGC |
| LsAP2-CRISPR/Cas9-R0 | AACCTTGGGACAGCGATCCTCCCAATCTCTTAGTCGACTCTAC |
| **Yeast two-hybrid assays** | |
| LsAP2-AD-F | CCAGATTACGCTCATATGATGTGGGATCTAAACGGGTTTC |
| LsAP2-AD-R | GGCCTCCATGGCCATATGTCAAACATTGAATCTTTGCTGT |
| LsAS1-BD-F | GAGGACCTGCATATGATGAAGGAGCGTCAACGATG |
| LsAS1-BD-R | CTCCATGGCCATATGTTAACTTAAACCATGATGATGATTAGTTCC |
| LsAS2-BD-F | GAGGACCTGCATATGATGGCGTCGTCTTCGAATTCG |
| LsAS2-BD-R | CTCCATGGCCATATGTTAAGATGGATCAATGGGTGTTCTCC |
| LsBP-BD-F | GAGGACCTGCATATGATGGAAGATTACATGGGCCAAAT |
| LsBP-BD-R | CTCCATGGCCATATGTCAAGGACCCAATCTGTAAGGAC |
| LsKAN1-BD-F | GAGGACCTGCATATGATGCCCTTGGAAGGGATTTTC |
| LsKAN1-BD-R | CTCCATGGCCATATGTCAGTTGCGCTCTTTTTGAAGC |

| | |
|---|---|
| LsKAN2-BD-F | GAGGACCTGCATATGATGGAGCTCTTTCCAGCACA |
| LsKAN2-BD-R | CTCCATGGCCATATGTCACCTTGAAGAATTGCTCCATAG |
| **LCI assays** | |
| LsAP2-cLUC-F | TCCCGGGGCGGTACCATGTGGGATCTAAACGGGTTTC |
| LsAP2-cLUC-R | GCTCTGCAGGTCGACTCAAACATTGAATCTTTGCTGT |
| LsAS2-nLUC-F | GACGAGCTCGGTACCATGGCGTCGTCTTCGAATTCG |
| LsAS2-nLUC-R | CGAGATCTGGTCGACAGATGGATCAATGGGTGTTC |
| LsBP-nLUC-F | GACGAGCTCGGTACCATGGAAGATTACATGGGCCAAAT |
| LsBP-nLUC-R | CGAGATCTGGTCGACAGGACCCAATCTGTAAGGAC |
| LsKAN1-nLUC-F | GACGAGCTCGGTACCATGCCCTTGGAAGGGATTTTC |
| LsKAN1-nLUC-R | CGAGATCTGGTCGACGTTGCGCTCTTTTTGAAGC |
| LsKAN2-nLUC-F | GACGAGCTCGGTACCATGGAGCTCTTTCCAGCACA |
| LsKAN2-nLUC-R | CGAGATCTGGTCGACCCTTGAAGAATTGCTCCATAG |
| **Dual-luciferase reporter assays** | |
| LsAP2-62-SK-F | AGAACTAGTGGATCCATGTGGGATCTAAACGGGTTTC |
| LsAP2-62-SK-R | GGTATCGATAAGCTTTCAAACATTGAATCTTTGCTGT |
| LsBP-62-SK-F | AGAACTAGTGGATCCATGGAAGATTACATGGGCCAAAT |
| LsBP-62-SK-R | GGTATCGATAAGCTTTCAAGGACCCAATCTGTAAGGAC |
| pBP-0800-LUC-F | GGTATCGATAAGCTTGTCATGTGAATACGTTCAACCTC |
| pBP-0800-LUC-R | AGAACTAGTGGATCCCACTCTTGCTCAAGAAAAACCCTAC |
| **Transgenic plants identification** | |
| CRISPR/Cas9-ID-F | TGTCCCAGGATTAGAATGATTAGGC |
| CRISPR/Cas9-ID-R | AGCCCTCTTCTTTCGATCCATCAAC |
| GUS-ID-F1 | GTGCATGGCTGGATATGTATCAC |
| GUS-ID-R1 | ACCCATCTCATAAATAACGTCATGC |
| GUS-ID-F2 | CAAGATCTAGTGGGAGATGCAC |
| GUS-ID-R2 | GACGTTGCCCGCATAATTACG |
| **Off-target sites detection** | |
| T1-OFF1-F | CTTGCCGTCAATACATGCAC |
| T1-OFF1-R | GTGGATAAACAATTATTGTGTAGACG |
| T1-OFF2-F | CAAGATACCACCACTATTGAGG |
| T1-OFF2-R | GTTTTCCTTTGTGGTCGGTC |
| T1-OFF3-F | CTAGACAATTGATCGCAATGC |
| T1-OFF3-R | CTGTAACAGAGATAGTCGCTCG |
| T1-OFF4-F | CTAGATGCAACCTGTTATTGTAC |
| T1-OFF4-R | GTAGATGTCAACTCAGATGATGC |
| T1-OFF5-F | CATGTAGGAATCACTTGTGTGTAC |
| T1-OFF5-R | GTTCCTCAACAGATGGCATC |
| T1-OFF6-F | TGTGCATCGATGGAGAGTATG |
| T1-OFF6-R | GCCAGCTACAAATTCTAACAC |
| T2-OFF1-F | GAGGATGACACGTGTCACAC |
| T2-OFF1-R | ACATTCCTAACACCAAATGGAC |
| T2-OFF2-F | AGGAAGAGAGTAGATCTGGAGAC |
| T2-OFF2-R | CCCTATAAGACGAGTTTGGCTC |

| | |
|---|---|
| T2-OFF3-F | TGGGCCTTACCCTAAAGTCC |
| T2-OFF3-R | TTAACCCTAGGTGGTTGGATC |

**Table S3. FPKM values of lettuce *AP2* genes at different seed development stages.**

| Gene name | S0 | S1 | S3 | S6 | S9 |
|---|---|---|---|---|---|
| *LsAP2* | 119.44 | 78.98 | 51.21 | 15.15 | 3.03 |
| *LsAP2-like1* | 58.99 | 50.40 | 31.68 | 7.92 | 2.28 |
| *LsAP2-like2* | 0.77 | 0.95 | 0.48 | 0.65 | 0.31 |

**Table S4. Editing efficiency and mutation types of CRISPR/Cas9-mediated editing of *LsAP2* in T0 generation plants.**

| No. of transgenic plants (T0 generation) | No. of plants with mutation | Mutation frequency (%) | Mutation type | | |
|---|---|---|---|---|---|
| | | | Monoallelic (%) | Biallelic (%) | Homozygous (%) |
| 10 | 6 | 60 | 0 | 40 | 20 |

**Table S5. Summary of the putative off-target sites analyses.**

| Targets | Putative off-target sites | | | | No. of plants examined | No. of plants mutated |
|---|---|---|---|---|---|---|
| | Name | Sequence | Locus | No. of mismatches | | |
| Target 1 | T1-OFF1 | GAGCACAACAGGTAATGGGGGG | ch3: -95127915 | 3 | 10 | 0 |
| | T1-OFF2 | AAGTATTAAAGGTGATGGGGGG | ch5: -201437556 | 3 | 10 | 0 |
| | T1-OFF3 | AAAAACTATAAGTGATGGGAGG | ch7: +180615736 | 4 | 10 | 0 |
| | T1-OFF4 | AAGCACTAGAGATGAAGGGTGG | ch2: -111314092 | 3 | 10 | 0 |
| | T1-OFF5 | AAGCACTAGAGATGAAGGGTGG | ch4: -153269803 | 3 | 10 | 0 |
| | T1-OFF6 | AAGCAATGCAGGTGATGGTTGG | ch2: +182546667 | 3 | 10 | 0 |
| Target 2 | T2-OFF1 | GGAGAATGGCTGACCCAAGTGG | ch6: -143598720 | 3 | 10 | 0 |
| | T2-OFF2 | GGAGAATGGACGTCCCAAGTGG | ch8: -297250967 | 4 | 10 | 0 |
| | T2-OFF3 | GAAGGATCGAGGTCCAAAGAGG | ch5: -50487265 | 4 | 10 | 0 |

**Data S1. Coding sequences and amino acid sequences of *LsAP2* gene in WT and three mutant lines.**

Start codons are highlighted in green, stop codons are highlighted in purple, the AP2 domains are highlighted in gray, target sequences are highlighted in yellow, PAM sequences are highlighted in red, and the insertion is highlighted in blue.

**WT**

**Coding sequence**

ATGTGGGATCTAAACGGGTTTCCTGATCGGAGGGGGTGCTCTTCACCAGTAGAAGGGGATGAAGAGGA
TAGTAACAATACTAATAAGGGTAAAGGGGTGGGGTCGATATCGAATTCAAGTTCCTCCGCCGTTGTTAT
GGAGGAAGATGGGGGCTCCGAGGAGGAGGAATCGGAGAGAGGGAGTCCAAGGAAACGAAGCACTAC
AGGTGATGGGAGGTTGTTTGGTTTTTCTATGACGGAAAACTCCTCTTGGGACAGCGATCCTCCAGTCAC
ACGACAGTTTTTCCCGGTTGATGATTCCGAGGTGGGACCCACGACCTCCGGCGGAGTTGGTGAAGGGT
TGATCTCACCGGCTACAACTTTCCCGACTGCTCACTGGTTTGGAGTTAAATTCTGCCACCAGTCATCCG
ATCCTTTAGATGGTGTCGCCGCCACAGCAGATGGGTTCCTCGGAAAAACTACCCCCGCCGTTGTTCCTC
AGCCGCTGAAGAAAAGCCGCCGTGGGCCGCGGTCTAGAAGTTCTCAGTACCGTGGAGTCACCTTTTAC
CGGCGAACCGGCCGATGGGAATCACATATATGGGATTGTGGAAAACAAGTTTATCTAGGTGGATTTGAT
ACTGCACACGCAGCGGCACGTGCATATGATAGGGCAGCGATTAAGTTTCGAGGAATGGAAGCCGATAT
AAACTTTAATCTTGAAGATTATGAAGACGATGTAAAACAGATGAGCACTCTAACAAAGGAAGAGTTTG
TGCATGTACTAAGGAGACAAAGTACTGGATTTCCAAGAGGGAGTTCAAAATACAGAGGTGTAACTTTG
CATAAATGTGGTAGATGGGAAGCTAGAATGGGCCAATTTTTAGGCAAAAAGTATGTGTATTTGGGTCTG
TTTGATACAGAAGTTGAAGCAGCCAGGGCATATGACAAGGCTGCTATCAAATGTAATGGAAAAGCCGC
TGTTACCAATTTTGATTCAAGTATATATGACAATGAGCTCAATTTGACTGAATGTTCAAACAATAGCAAT
AAAGATAGCAATGTATCATCGGACCATAATCTTGATTTGAGCCTAGGTGGTGGCTCGGCTTCAAATACGA
TGAATGAGGGAGGAGAAAATGGTCATTTTACAAGGGATCATCATCATTCTGACACCATCCAACTTGGGT
TTAGACCTAGTGGACGCAATGAGACAAGGACATCGCATTTTATGGACCATAGAAATCTTGAATCTTTTAA
TATTCCGAATGAAATACGTGGATACAAACATTTTATGAGACCCGTTGACTCGTCTATGCATAATATGTTCG
ATCCACCAATCTTCAACTCATTGAGTCATCAAATGCAGTTTTCCAGCAGCACAATGCAAGGAGGAAATA
ATTTATCTGCAGCTTCAATTGGTGAAAATAATATGAATGCCAATGCACCTCCTCTACACCAAATCTATGC
AAAAAGTGCTGCAGCATCATCAGGATTCCCACAGCAAAGATTCAATGTTTGA

**Amino acid sequence**

MWDLNGFPDRRGCSSPVEGDEEDSNNTNKGKGVGSISNSSSSAVVMEEDGGSEEEESERGSPRKRSTTGD
GRLFGFSMTENSSWDSDPPVTRQFFPVDDSEVGPTTSGGVGEGLISPATTFPTAHWFGVKFCHQSSDPLDGV
AATADGFLGKTTPAVVPQPLKKSRRGPRSRSSQYRGVTFYRRTGRWESHIWDCGKQVYLGGFDTAHAAAR
AYDRAAIKFRGMEADINFNLEDYEDDVKQMSTLTKEEFVHVLRRQSTGFPRGSSKYRGVTLHKCGRWEA
RMGQFLGKKYVYLGLFDTEVEAARAYDKAAIKCNGKAAVTNFDSSIYDNELNLTECSNNSNKDSNVSSD
HNLDLSLGGGSASNTMNEGGENGHFTRDHHHSDTIQLGFRPSGRNETRTSHFMDHRNLESFNIPNEIRGYK
HFMRPVDSSMHNMFDPPIFNSLSHQMQFSSSTMQGGNNLSAASIGENNMNANAPPLHQIYAKSAAASSGF
PQQRFNV

**#30-A1**

**Coding sequence**

ATGTGGGATCTAAACGGGTTTCCTGATCGGAGGGGGTGCTCTTCACCAGTAGAAGGGGATGAAGAGGA
TAGTAACAATACTAATAAGGGTAAAGGGGTGGGGTCGATATCGAATTCAAGTTCCTCCGCCGTTGTTAT
GGAGGAAGATGGGGGCTCCGAGGAGGAGGAATCGGAGAGAGGGAGTCCAAGGAAACGAAGCACTAC
AGGTGATGGGACAGCGATCCTCCAGTCACACGACAGTTTTTCCCGGTTGATGATTCCGAGGTGGGACC
CACGACCTCCGGCGGAGTTGGTGAAGGGTTGA

**Amino acid sequence**

MWDLNGFPDRRGCSSPVEGDEEDSNNTNKGKGVGSISNSSSSAVVMEEDGGSEEEESERGSPRKRSTTGD
GTAILQSHDSFSRLMIPRWDPRPPAELVKG

**#49**

**Coding sequence**

ATGTGGGATCTAAACGGGTTTCCTGATCGGAGGGGGTGCTCTTCACCAGTAGAAGGGGATGAAGAGGA
TAGTAACAATACTAATAAGGGTAAAGGGGTGGGGTCGATATCGAATTCAAGTTCCTCCGCCGTTGTTAT
GGAGGAAGATGGGGGCTCCGAGGAGGAGGAATCGGAGAGAGGGAGTCCAAGGAAACGAAGCACTAC
AGGTGATTGGGAGGTTGTTTGGTTTTTCTATGACGGAAAACTCCTCTTGGGACAGCGATCCTCCAGTCA
CACGACAGTTTTTCCCGGTTGA

**Amino acid sequence**

MWDLNGFPDRRGCSSPVEGDEEDSNNTNKGKGVGSISNSSSSAVVMEEDGGSEEEESERGSPRKRSTTGD
WEVVWFFYDGKLLLGQRSSSHTTVFPG

**#100-A1**

**Coding sequence**

ATGTGGGATCTAAACGGGTTTCCTGATCGGAGGGGGTGCTCTTCACCAGTAGAAGGGGATGAAGAGGA
TAGTAACAATACTAATAAGGGTAAAGGGGTGGGGTCGATATCGAATTCAAGTTCCTCCGCCGTTGTTAT
GGAGGAAGATGGGGGCTCCGAGGAGGAGGAATCGGAGAGAGGGAGTCCAAGGAAACGAAGCACTAC
ATGGGAGGTTGTTTGGTTTTTCTATGACGGAAAACTCCTCTTGGGACAGCGATCCTCCAGTCACACGAC
AGTTTTTCCCGGTTGA

**Amino acid sequence**

MWDLNGFPDRRGCSSPVEGDEEDSNNTNKGKGVGSISNSSSSAVVMEEDGGSEEEESERGSPRKRSTTWE
VVWFFYDGKLLLGQRSSSHTTVFPG
